# Convergent evolution of SARS-CoV-2 spike mutations, L452R, E484Q and P681R, in the second wave of COVID-19 in Maharashtra, India

**DOI:** 10.1101/2021.04.22.440932

**Authors:** Sarah Cherian, Varsha Potdar, Santosh Jadhav, Pragya Yadav, Nivedita Gupta, Mousmi Das, Partha Rakshit, Sujeet Singh, Priya Abraham, Samiran Panda, NIC team

**Affiliations:** ICMR-National Institute of Virology, Pune; Indian Council of Medical Research, N. Delhi; National Centre for Disease Control, New Delhi

## Abstract

As the global severe acute respiratory syndrome coronavirus 2 (SARS-CoV-2) pandemic expands, genomic epidemiology and whole genome sequencing are being constantly used to investigate its transmissions and evolution. In the backdrop of the global emergence of “variants of concern” (VOCs) during December 2020 and an upsurge in a state in the western part of India since January 2021, whole genome sequencing and analysis of spike protein mutations using sequence and structural approaches was undertaken to identify possible new variants and gauge the fitness of current circulating strains.

Phylogenetic analysis revealed that the predominant clade in circulation was a distinct newly identified lineage B.1.617 possessing common signature mutations D111D, G142D, L452R, E484Q, D614G and P681R, in the spike protein including within the receptor binding domain (RBD). Of these, the mutations at residue positions 452, 484 and 681 have been reported in other globally circulating lineages. The structural analysis of RBD mutations L452R and E484Q along with P681R in the furin cleavage site, revealed that these may possibly result in increased ACE2 binding and rate of S1-S2 cleavage resulting in better transmissibility. The same two RBD mutations indicated decreased binding to select monoclonal antibodies (mAbs) and may affect their neutralization potential. Experimental validation against a wider panel of mAbs, sera from vaccinees and those that recovered from natural infection needs to be studied.

The emergence of such local variants through the accumulation of convergent mutations during the COVID-19 second wave needs to be further investigated for their public health impact in the rest of the country and its possibility of becoming a VOC.

## Introduction

The severe acute respiratory syndrome coronavirus 2 (SARS-CoV-2) surface spike (S) protein mediates entry into host cells by binding to the host receptor angiotensin-converting enzyme 2 (ACE2) via its receptor-binding domain (RBD). Crystal structures of SARS-CoV-2 S protein or its RBD complexed with ACE2 from different hosts reveal that the RBD contains a core and a receptor-binding motif (RBM) which forms contacts with ACE2^1^. A number of naturally selected mutations in the RBM have been shown to affect infectivity, human-to-human transmission, pathogenesis and immune escape^2^.

As the global SARS-CoV-2 pandemic expands, genomic epidemiology and whole genome sequencing are being constantly used to investigate the transmissions and evolution. The expansion of the genome-wide diversity resulted in delineation of the viral strains into clades, lineages, and sub-lineages. Clades G/GH/GR/GV/GRY as per the Global Initiative on Sharing All Influenza Data (GISAID) database (https://www.gisaid.org/)^3^ possess a common mutation D614G in the spike protein. The mutation was shown to result in significantly higher human host infectivity and better transmission efficiency to the virus^4,5^. Three recently emerged “variants of concern” (VOCs) of SARS-CoV-2 are the UK-variant 501Y.V1/B.1.1.7, South Africa variant 501.V2/ B.1.351 and Brazilian variant 501Y.V3/P.1 (alias of lineage B.1.1.28.1). These variants are known to possess multiple mutations across the genome, including several in the S protein and its RBD such as N501Y, E484K and K417N/T^5–7^. Multiple SARS-CoV-2 variants are now seen to be circulating globally.

The potential introduction and consequence of these emerging variants in the country is vital to support the public health response. The National Influenza Centre at the ICMR-National Institute of Virology, Pune, as an apex laboratory of the Indian Council of Medical Research (ICMR) has been continuously involved in the SARS-CoV-2 diagnostics and monitoring the genomic evolution of this virus. In addition, the ICMR-NIV, Pune is also one among the ten national laboratories of the Indian SARS-CoV-2 Consortium on Genomics (INSACOG), Ministry of Health and Family Welfare, Government of India, catering to western India. In the backdrop of the global emergence of VOCs and an upsurge in Maharashtra, a state in the western part of India, enhanced sequencing was undertaken. This was to identify possible new variants and analyze spike protein mutants, in particular, to gauge the fitness of current circulating strains, using bioinformatics sequence and molecular structural approaches.

## Materials and methods

The State of Maharashtra (latitude 19°39’ N, longitude 75°18’E) is located in west central India, in the north western part of the Indian Subcontinent. Since the end of January 2021, a concentrated spurt in COVID-19 cases was noted in several districts of Maharashtra (**Suppl. Fig. 1**). As per the State Government directives, samples from international travelers and 5% of surveillance samples from the positive cases with Ct value <25 including those from clusters, long haulers, mild/moderate/severe and deceased cases, were referred to the ICMR-NIV, Pune for whole genome sequencing. Nasopharyngeal swabs (n=733, November 25, 2020-March 31, 2021) were processed for whole genome sequencing.

Briefly, nucleic acid extraction was performed using 280 μL of each sample in duplicate by Qiagen viral RNA extraction protocol and quantified RNA was further processed for template preparation using the Ion Chef System. Purified template beads were submitted to meta transcriptome next-generation sequencing (NGS) in the Ion S5 platform (ThermoFisher Scientific) using an Ion 540™ chip and the Ion Total RNA-Seq kit v2.0, as per the manufacturer’s protocol (ThermoFisher Scientific)^8^. Sequence data was processed using the Torrent Suite Software (TSS) v5.10.1 (ThermoFisher Scientific, USA). Coverage analysis plugins were utilized to generate coverage analysis report for each of the samples. Reference-based reads gathering, and assembly were performed for all the samples using Iterative refinement meta-assembler (IRMA)^9^. A subset of the samples (were also sequenced on the Illumina machine and analyzed using on the CLC Genomics Workbench version 20 (CLC, Qiagen) as described elsewhere^10^. Of 733 whole genomes obtained, 598 were considered for different analyses based on coverage of ≥ 93% of the genome. For each whole genome sequence, the GISAID Clade assignment, was done using CoVsurver: Mutation Analysis of hCoV-19 (https://www.gisaid.org/epiflu-applications/covsurver-mutations-app/). For lineage assignment, the web application ie., Phylogenetic Assignment of Named Global Outbreak LINeages (PangoLIN) COVID-19 Lineage Assigner (https://pangolin.cog-uk.io/), was implemented^7^. The 598 genomes were aligned using MUSCLE and a Maximum Likelihood phylogenetic tree was constructed using IQ-tree v1.6^11,12^, employing the GTR as the substitution model and 1,000 bootstrap replications. The average frequency of nonsynonymous mutations in the genomes over the period from 1st December, 2020 to 31^st^ March, 2021 was calculated, considering Wuhan as the reference strain.

For further structural characterization of the S protein mutations, the crystal structure of SARS-CoV-2 S glycoprotein complexed with ACE2 was obtained from the protein data bank (PDB ID: 7A98^13^. The top ten mutations in the S protein were mapped using Biovia Discovery studio visualizer 2020. To assess the effect of noted RBD mutations on ACE2 binding, the crystal structure of SARS-CoV-2 spike RBD domain complexed with ACE2 (PDB ID: 6LZG^14^) was used. For assessment of the noted mutations on binding to neutralizing antibodies, the SARS-CoV-2 spike RBD domain complexed with two selected mAbs REGN10933/ P2B-2F6 were retrieved (PDB ID: 6XDG; resolution 3.90Å and 7BWJ; resolution 2.65 Å respectively)^15,16^. Point mutations were carried out using Biovia Discovery studio visualizer 2020 and the structures of the complexes were subjected to energy minimization using macro model tool in Schrodinger 2020 using default parameters. The molecular interactions between the RBD-ACE2 interface, within the RBD and between the neutralizing mAbs-RBD were analyzed using non-bonded interactions tool in Biovia Discovery studio visualizer 2020.

## Results

The genomic surveillance for the spurt in the COVID-19 cases in Maharashtra was carried out to identify the circulating lineages/VOCs and identify possible functionally-significant mutations in the spike protein.

PangoLIN lineage classification of 598 whole genomes revealed the presence of 47 lineages (**Supplementary Table 1**), all within GISAID clade G, evident from the D614G mutation in the S protein. Lineages B.1.617 (n=273), B.1.36.29 (n=73), B.1.1.306 (n=67), B.1.1.7 (n=31), B.1.1.216 (n=24), B.1.596 (n=17), B.1 (n=17), B.1.1 (n=15) and B.1.36 (n=12) were found to be the predominant lineages (**Supplementary Figure 1**). Phylogenetic analysis revealed four sub-clusters within B.1.617 that could be linked to mutations specific to the spike region (**Figure 1**). Among these, cluster B.1.617:C (n=107), included majority of the strains from eastern part of Maharashtra while B.1.617:B (n=80) also included sequences from major cities like Pune, Thane and Aurangabad, more on the western part of the State. The frequency of mutations L452R and E484Q within the RBD and mutations G142D and P681R within the spike but outside the RBD region, increased from January 2021 (**Figure 2**). The P681R is noted in the S1-S2 furin-cleavage site. A mutation H1101D specific to clusters B.1.617: A and B.1.617:B, was not noted in January 2021 (**Figure 1**). The B.1.617:C cluster possessed a T95I-specific mutation. Notably a small cluster of B.1.617 (n=21) emerged in late March and did not possess the E484Q mutation. This clustered with lineage B.1.596 and shared common mutations T19R and D950N in the spike protein. A synonymous mutation D111D was observed to be co-occurring with the RBD mutations L452R and E484Q and it was absent in the cluster that did not possess E484Q (Figure 1). Based on the monthly averages of non-synonymous mutation events when compared to the Wuhan reference strain NC_045512.2, an increased frequency of non-synonymous mutations was noted in February 2021 (**Figure 3**).

**Figure 1:**
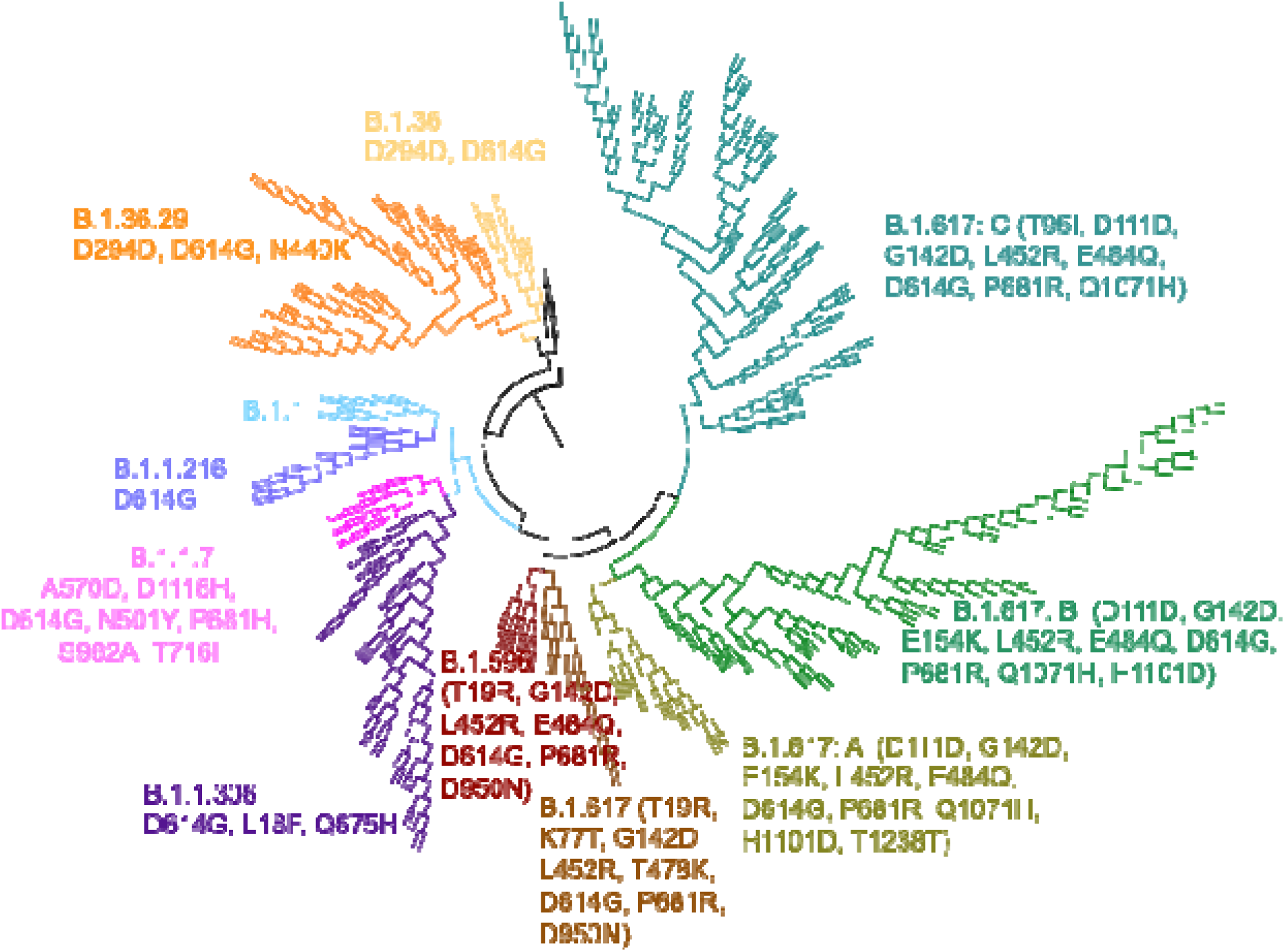
Maximum-likelihood tree of representative SARS-CoV-2 genomes depicting PangoLIN lineages and the co-occurring mutations in the spike protein in the sub-clusters of lineage B.1.617.

**Figure 2:**
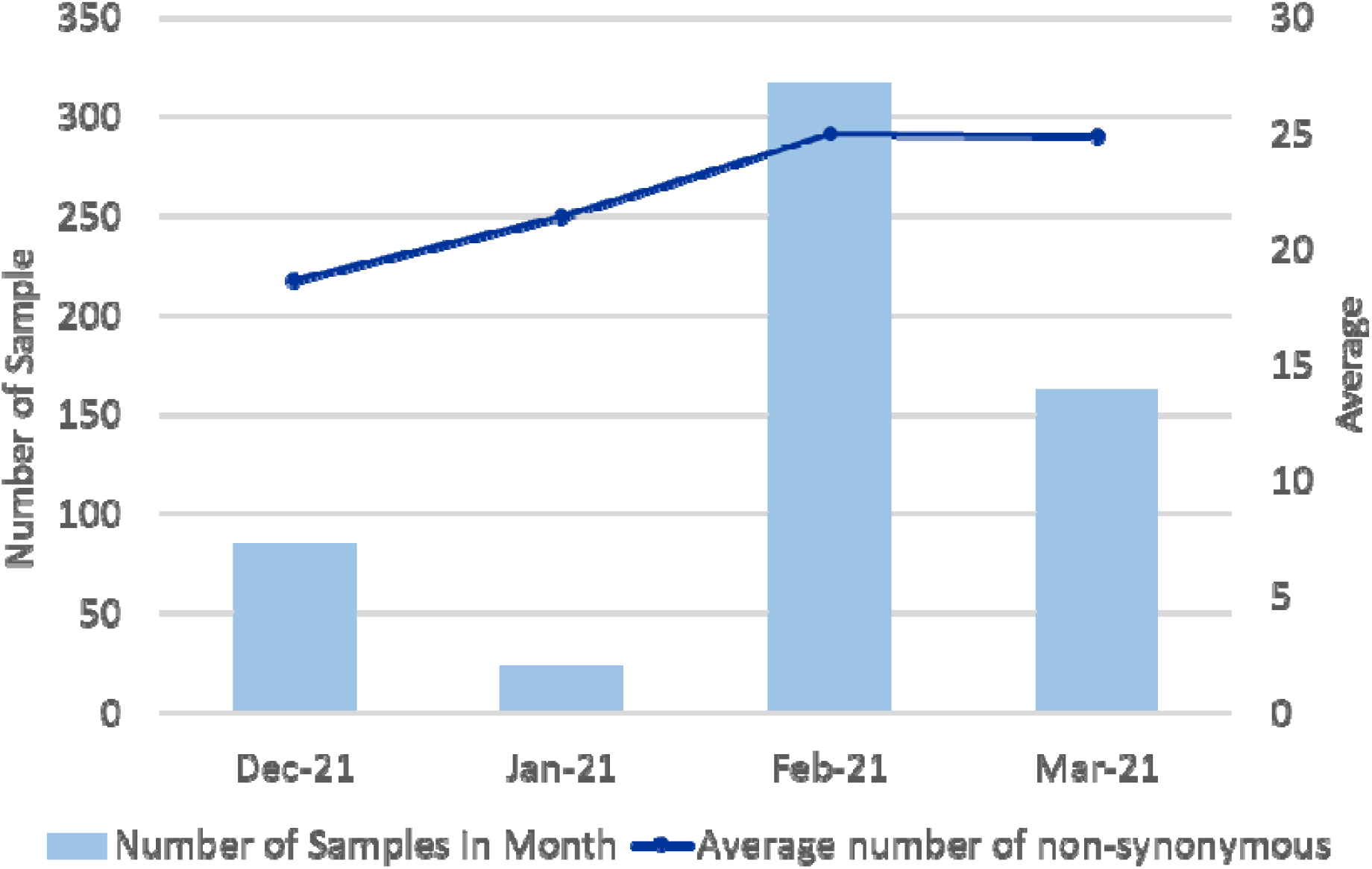
Monthly average of non-synonymous mutations (trend-line) against the total SARS-CoV-2 samples studied (blue bars).

**Figure 3:**
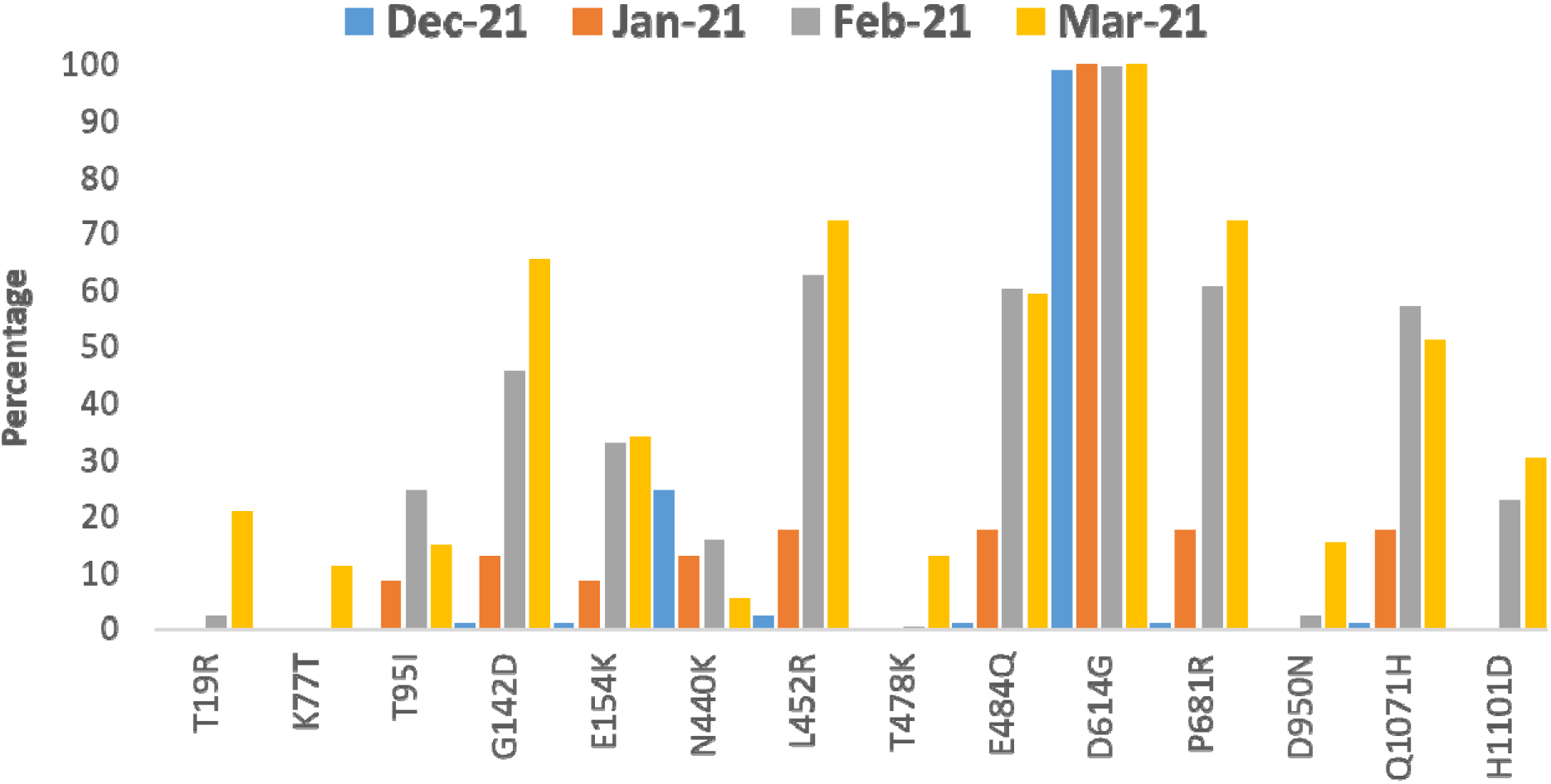
Trend of major mutations in the spike protein from December, 2020 to March, 2021.

The key mutations in the S protein are mapped on a furin-cleaved structure of the S protein (**Figure 4**). The structural implications of the RBD mutations (L452R and E484Q) were analyzed in terms of interaction with the ACE2 receptor and neutralizing antibodies that are known to have interactions with these residues. Effect of the E484Q mutation is noted in terms of disruption in an electrostatic bond of the spike RBD residue E484 with K31 in the ACE2 interaction interface (**Figure 5A, Supplementary Table 2**). The intramolecular interactions in the wild strain indicate that the L452 residue is involved in a hydrophobic interaction with L492 which forms another hydrophobic contact with F490. These residue interactions form a hydrophobic patch on the surface of the RBD. The L452R mutation abolishes the hydrophobic interaction with L492 of the RBD. Estimation of the minimized energies of the wild and mutant structures of RBD complexed with ACE2 showed energy values of −93732.305 kcal/mol and −94543.180 kcal/mol respectively.

**Figure 4:**
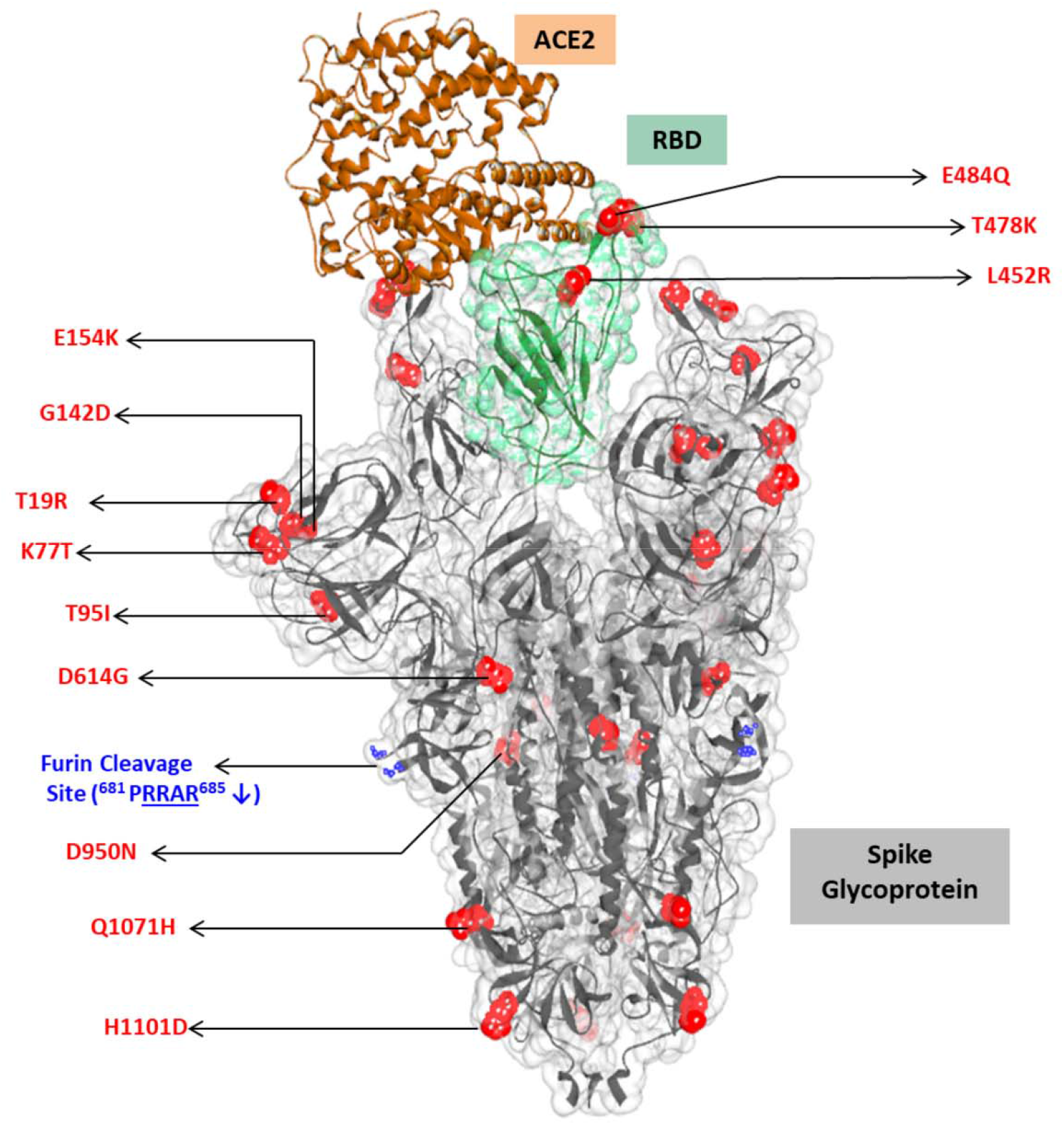
Mapping of key mutations on the furin-cleaved crystal structure of SARS-CoV-2 spike glycoprotein (grey surface view) in complex with ACE2 (brown solid ribbon). RBD region shown in green

**Figure 5:**
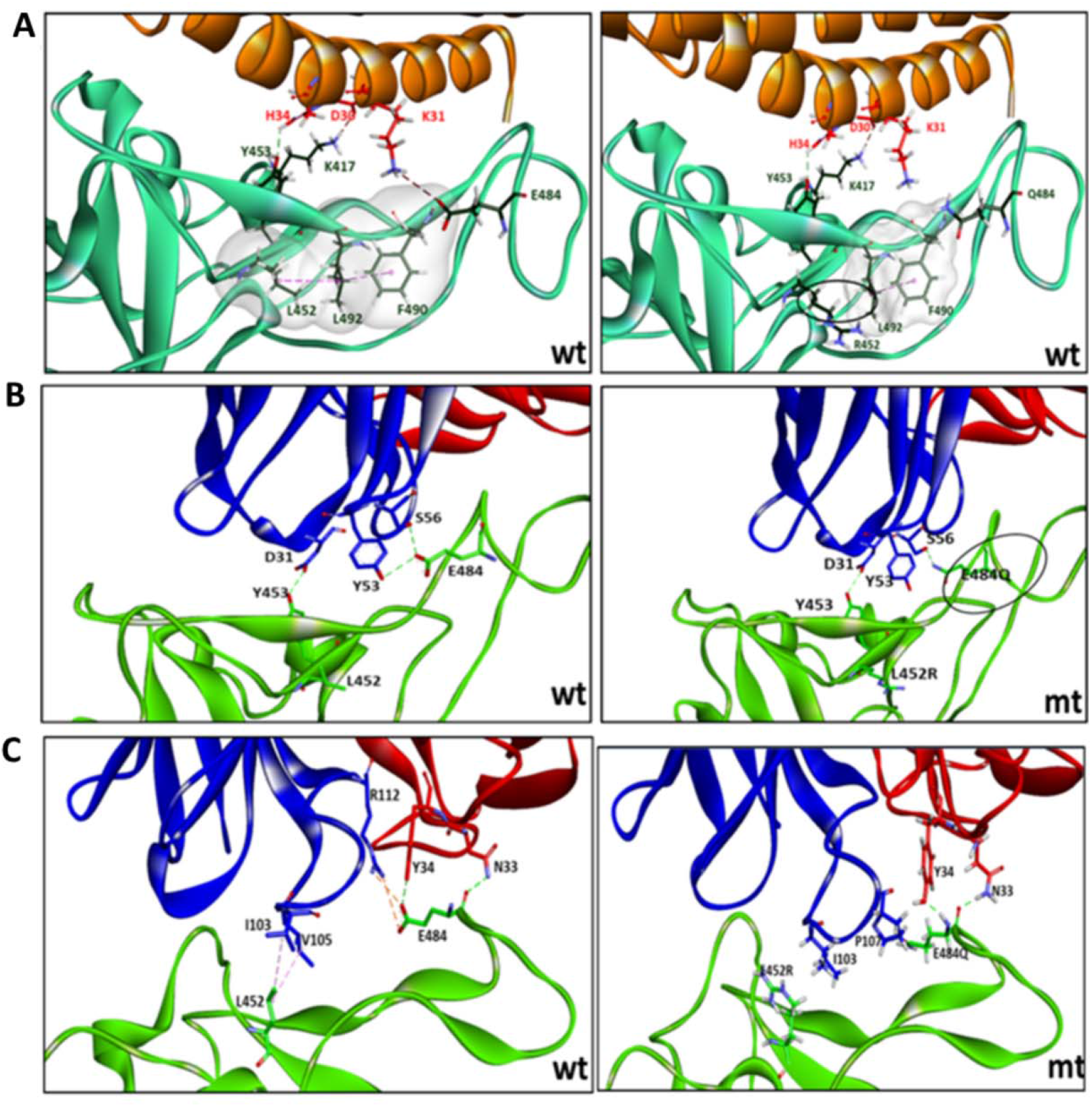
(**A**) Key interactions between ACE2-RBD involving major mutational hotspots in the RBD (**B**) Interactions between RBD-mAb REGN10933 (**C)** Interactions between RBD-mAb P2B-2F6. wt, corresponds to wild-type strain and mt, mutant strain. In (A), also shown are the intra-molecular contacts in a hydrophobic patch of the RBD region (surface displayed in grey color). In (B) and (C) blue represents heavy chain and red, light chain.

The mutations L452R and E484Q are seen to disrupt the interfacial interactions of spike RBD with specific neutralizing antibodies (**Figure 5B, C; Supplementary Table 3**), The heavy chain of monoclonal antibody REGN10933 interacts with the RBD by making two hydrogen bonds (H-bonds) between E484 of the RBD and Y53 and S56 of the antibody. Residue Y453 of the RBD makes an H-bond with D31 of the heavy chain of the antibody. The RBD mutation E484Q disrupts the 2 H-bonds with S56 and Y53 (**Figure 5B**). Residues L452 and E484 possess direct contacts with the mAb P2B-2F6 (**Figure 5C**). The intermolecular interactions of the wild-type complex indicate that L452 is involved in hydrophobic interactions with residues I103 and V105 of the heavy chain of the antibody. Residue E484 forms H-bonds with N33 and Y34 of the light chain as well as a salt bridge and H-bond interaction with R112 side chain of the light chain. The L452R mutation breaks the hydrophobic interactions with both residues I103 and V105 and also disrupts the H-bond and electrostatic interaction with R112.

## Discussion

In addition to the three global VOCs, some of the global variants of interest (VUIs) include B.1.525/ VUI-202102/03 that emerged in UK (https://cov-lineages.org/global_report_B.1.525.html), B.1.1.49 that emerged in Denmark (https://cov-lineages.org/lineages/lineage_B.1.1.49.html) and A.23.1 that was first detected in Uganda (https://cov-lineages.org/global_report_A.23.1.html). The potential consequences of emerging variants are increased transmissibility, increased pathogenicity and the ability to escape natural or vaccine-induced immunity^17,18^. With this background, we sequenced and analyzed the whole genomes of SARS-CoV-2 from different districts of Maharashtra where a surge in COVID-19 activity was noted since the end of January 2021 after a gap of almost four months.

Phylogenetic and sequence analyses revealed that only a small proportion of sequences majorly in the month of December 2020, were detected as the VOC B.1.1.7, UK-variant. On the other hand, the B.1.617 lineage possessing common signature mutations G142D, L452R, E484Q, D614G and P681R, in the spike protein could be linked to the increase in the proportion of non-synonymous mutations in February 2021. In addition, districts in western Maharashtra, such as Pune, Mumbai, Thane and Nashik showed presence of multiple lineages in circulation in comparison to the dominance of lineage B.1.617 in eastern Maharashtra (Supplementary figure 1). The co-occurrence of synonymous mutations with the non-synonymous mutations observed is interesting and being reported^19,20^. Notably D111D was found to be associated with the signature mutations.

The structural analysis of the effect of RBD mutations L452R and E484Q towards ACE2 binding, revealed decrease in intra-molecular and intermolecular contacts with respect to the wild-type. However, the hydrophobic L452 residue mutation to the hydrophilic 452R might help in interaction with water molecules and overall stabilization of the complex, as was reflected in the lower minimum energy of the mutant complex. Another significant mutation P681R in the furin cleavage site, resulting in enhancement of the basicity of the poly-basic stretch, might help in increased rate of membrane fusion, internalization and thus better transmissibility. In vitro and/or *in vivo* studies to understand the phenotypic effects of the mutant strains would be vital in this regard. Structural analysis further showed that the two RBD mutations L452R and E484Q may decrease the binding ability of REGN10933 and P2B-2F6 antibodies to the variant strains, compared to that in the wild-type strain. A recent report^21^ revealed that the L452R mutation reduced or abolished neutralizing activity of 14 out of 35 RBD-specific mAbs including three clinical stage mAbs. Another recent study^22^ has demonstrated that the mutation L452R can escape from human leukocyte antigen (HLA)-24-restricted cellular immunity and can also increase viral infectivity, potentially promoting viral replication. The combination of the two RBD mutations noted in this study could affect the neutralization of the select mAbs. The neutralizing potential of a wider range of mAbs against the strains of lineage B.1.617 needs to be assessed. Further the level of neutralization efficacy of sera from vaccinees of different vaccine formulations and those who experienced natural infection also needs to be studied.

Mutations at both the residue positions 452 and 484 individually have been reported earlier. L452R has been noted in California lineages B.1.427 and B.1.429 while E484K mutation is common to the three VOCs having global impact. E484Q has also been reported in several sequences in the GISAID with the earliest strain noted in Denmark. P681H is one of the mutations in the UK-variant B.1.1.7 while P681R is one of the mutations in the VUI lineage A.23.1. The unique combination of mutations L452R, E484Q and P681R noted in this study is an indication of convergent evolution.

To summarize, the study investigated the S protein mutations associated with the COVID-19 cases in Maharashtra observed since the month of February 2021. The continuous increase in positivity could be attributed to signature mutations in the spike protein and functionally significant co-occurring mutations. As per GISAID submissions, the B.1.617 lineage has been reported from several countries including UK, USA, Switzerland, Germany, Singapore etc. However, the emergence of such local variants during the COVID-19 second wave in India needs to be further investigated for their public health impact and its possibility of becoming a VOC.

## Acknowledgement

Authors are grateful to the Director Health Services of Maharashtra and State Integrated Disease Surveillance Program (IDSP) for coordination of samples. We also acknowledge the cohesive efforts of NIC staff. The authors also acknowledge the support of Dr. Balram Bhargava, Secretary, Department of Health Research, Government of India and Director General, Indian Council of Medical Research, N. Delhi.

## Supplementary Material

**Figure S1:**
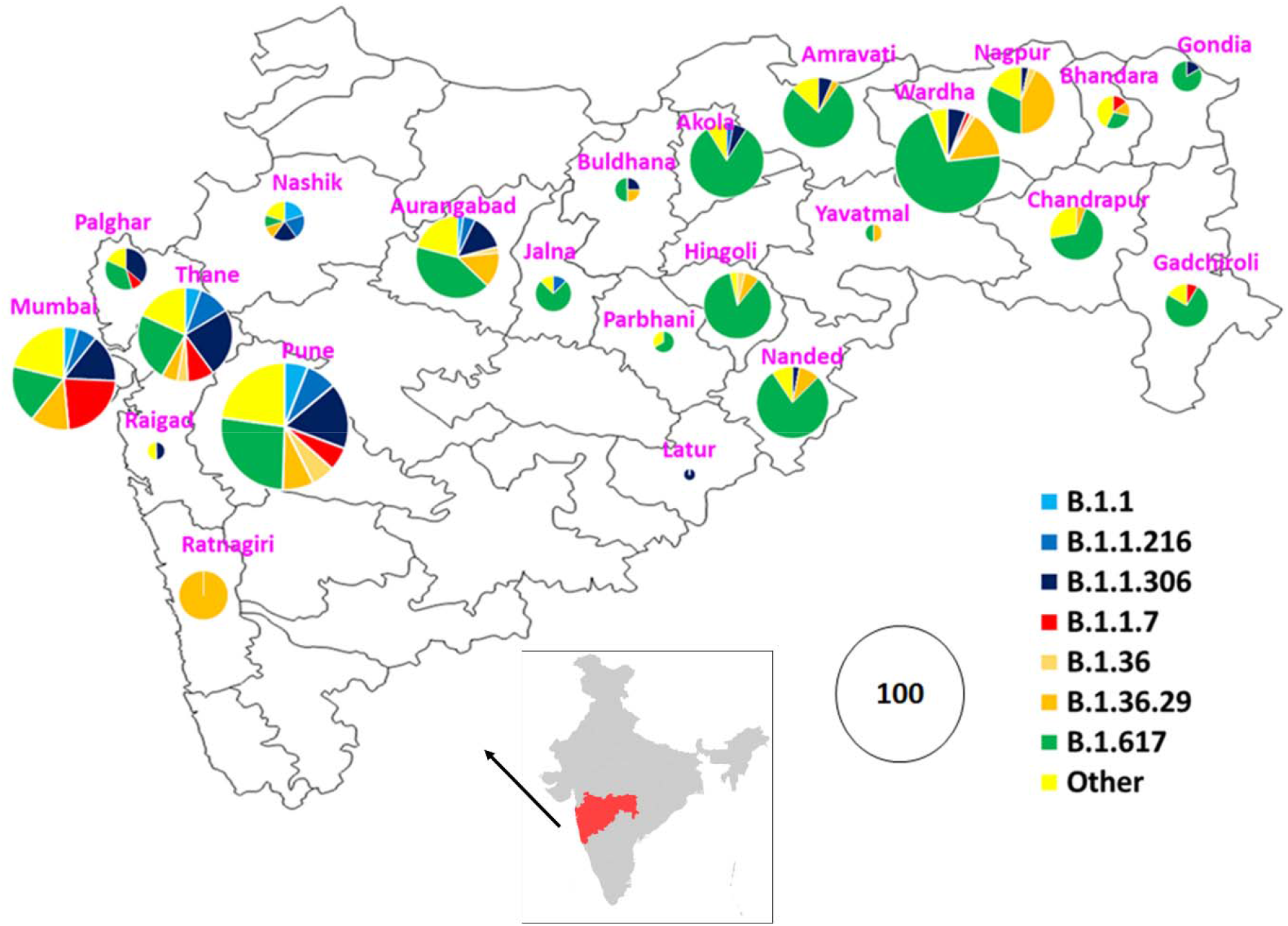
The distribution of predominant PangoLIN lineages in the labelled districts of Maharashtra. The size of the pie charts is proportional to the number of genome sequences from the particular district.

**Figure S2:**
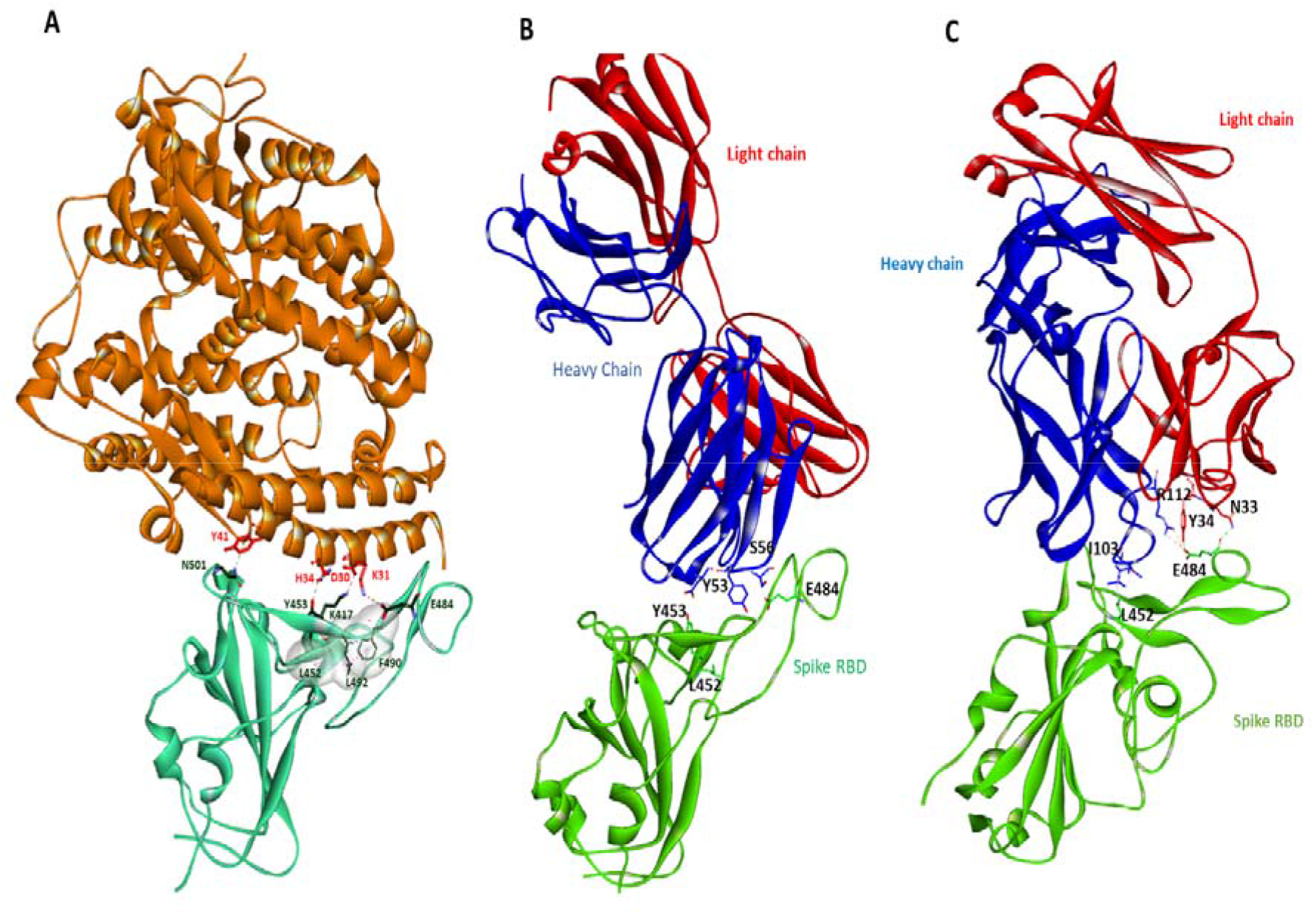
Structural complexes of RBD-ACE2 and RBD-Fab. **A**. RBD - ACE2 (PDB Id-6LZG), **B**. RBD - mAb REGN10933(PDB Id-6XDG) **C**. RBD - mAb P2B-2F6(7bwj)

**Supplementary Table 1:**
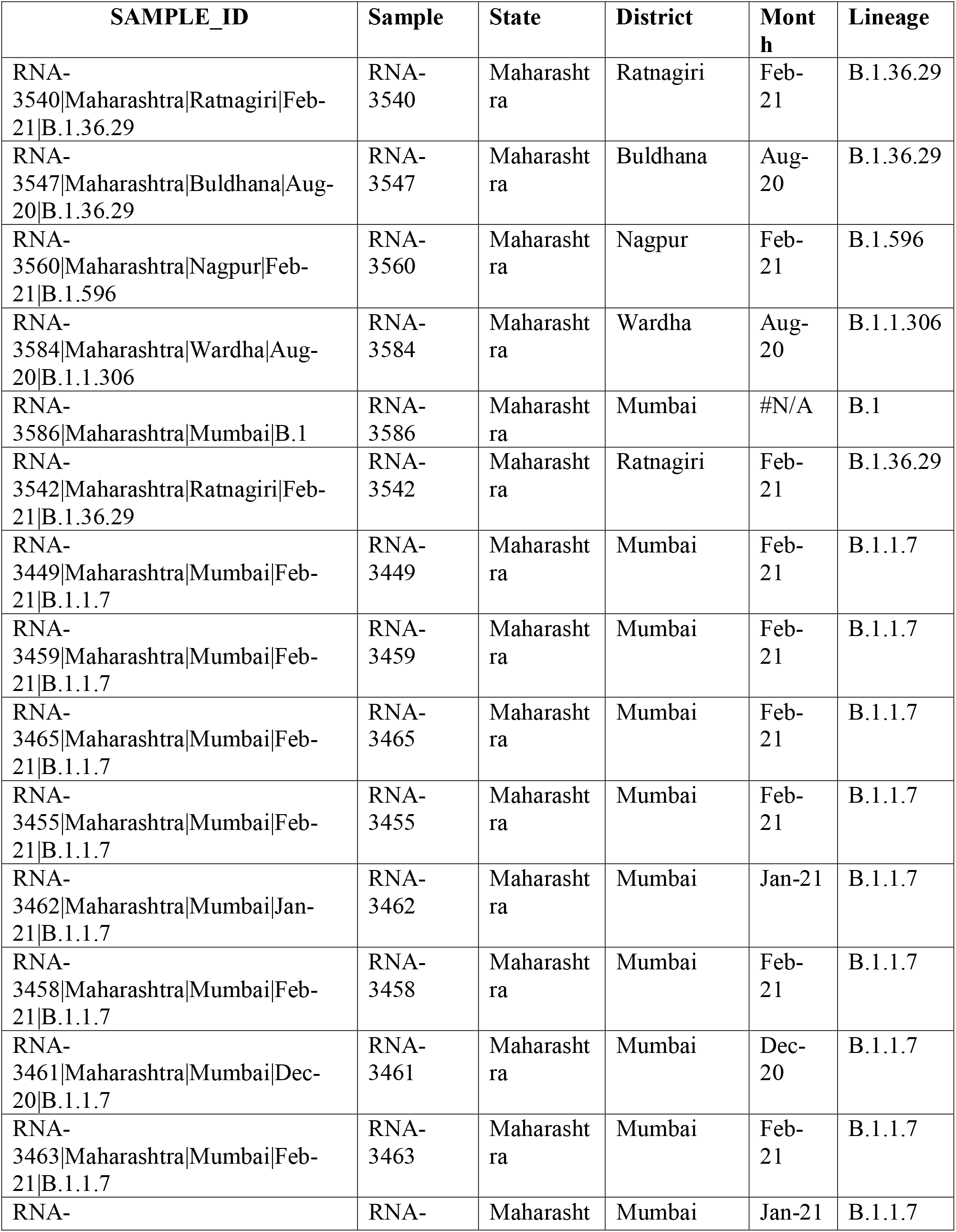
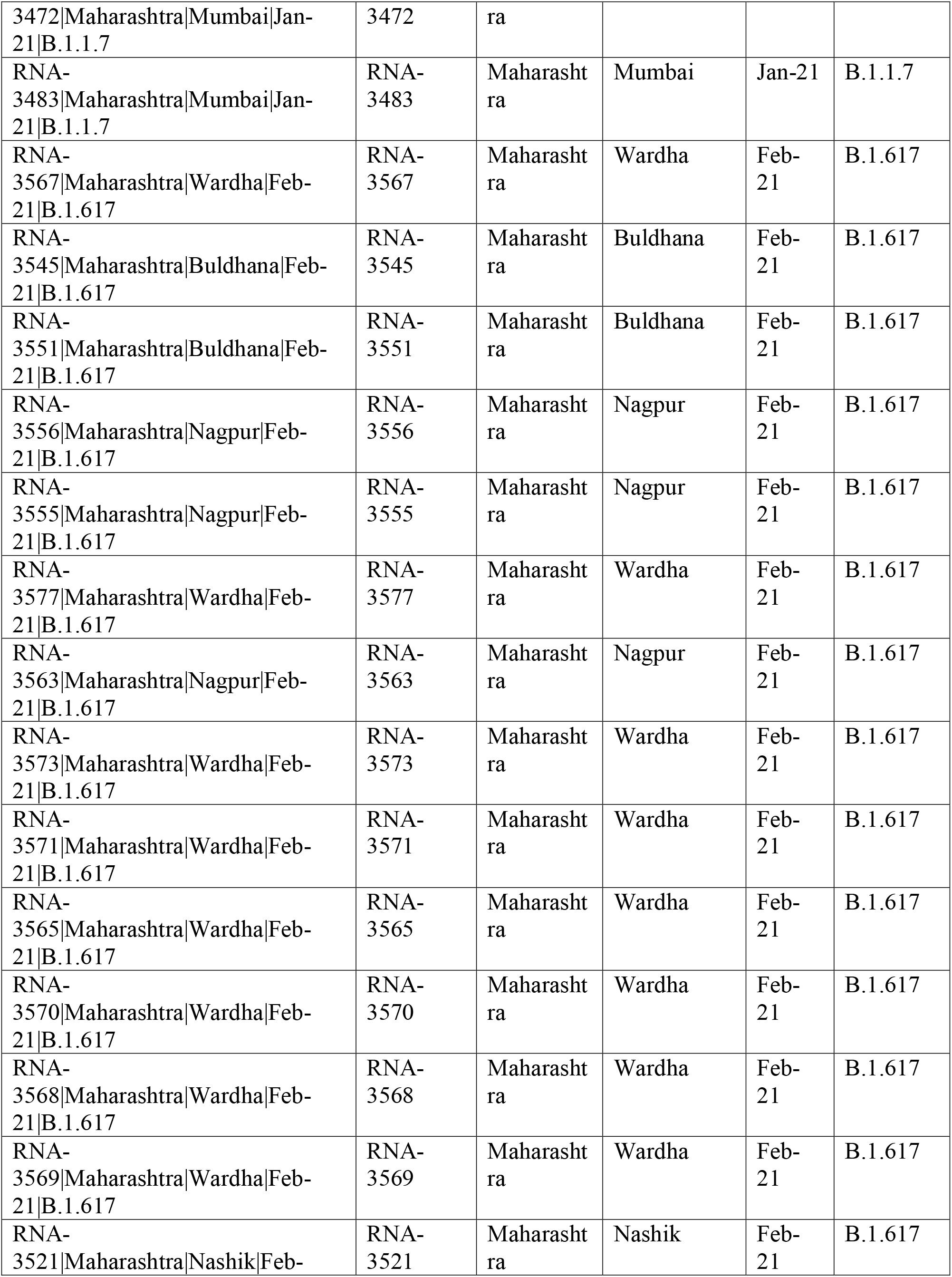
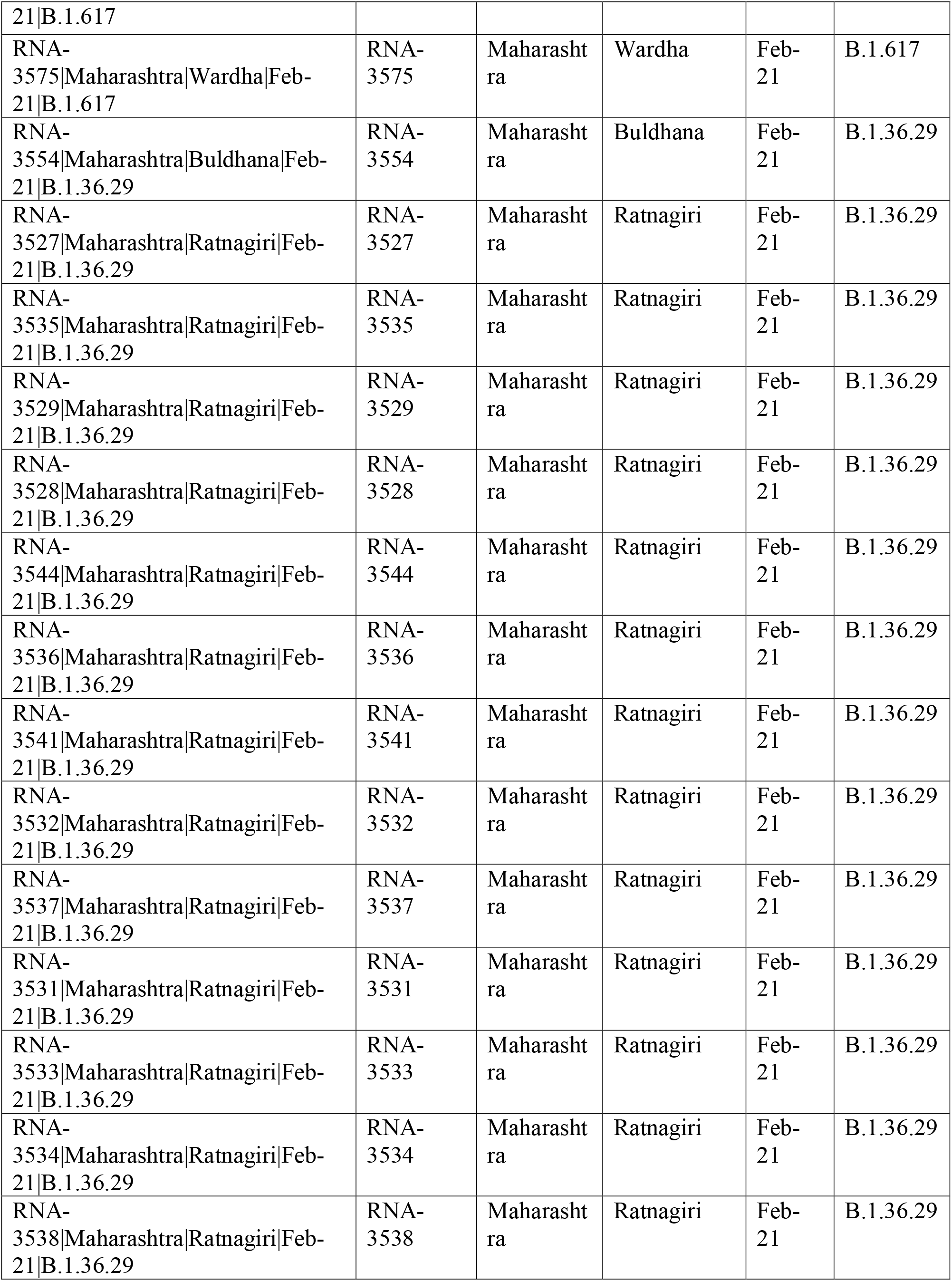
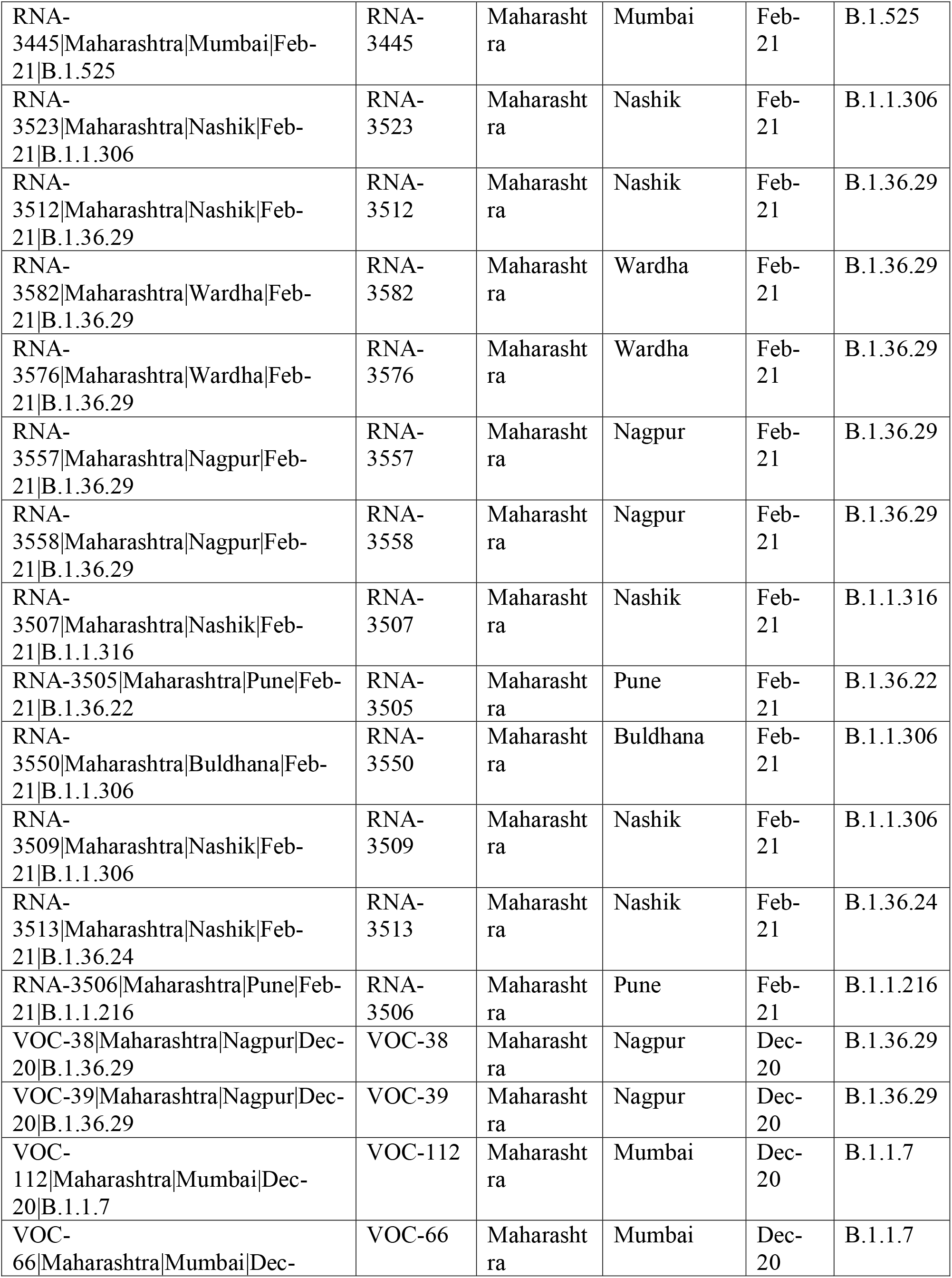
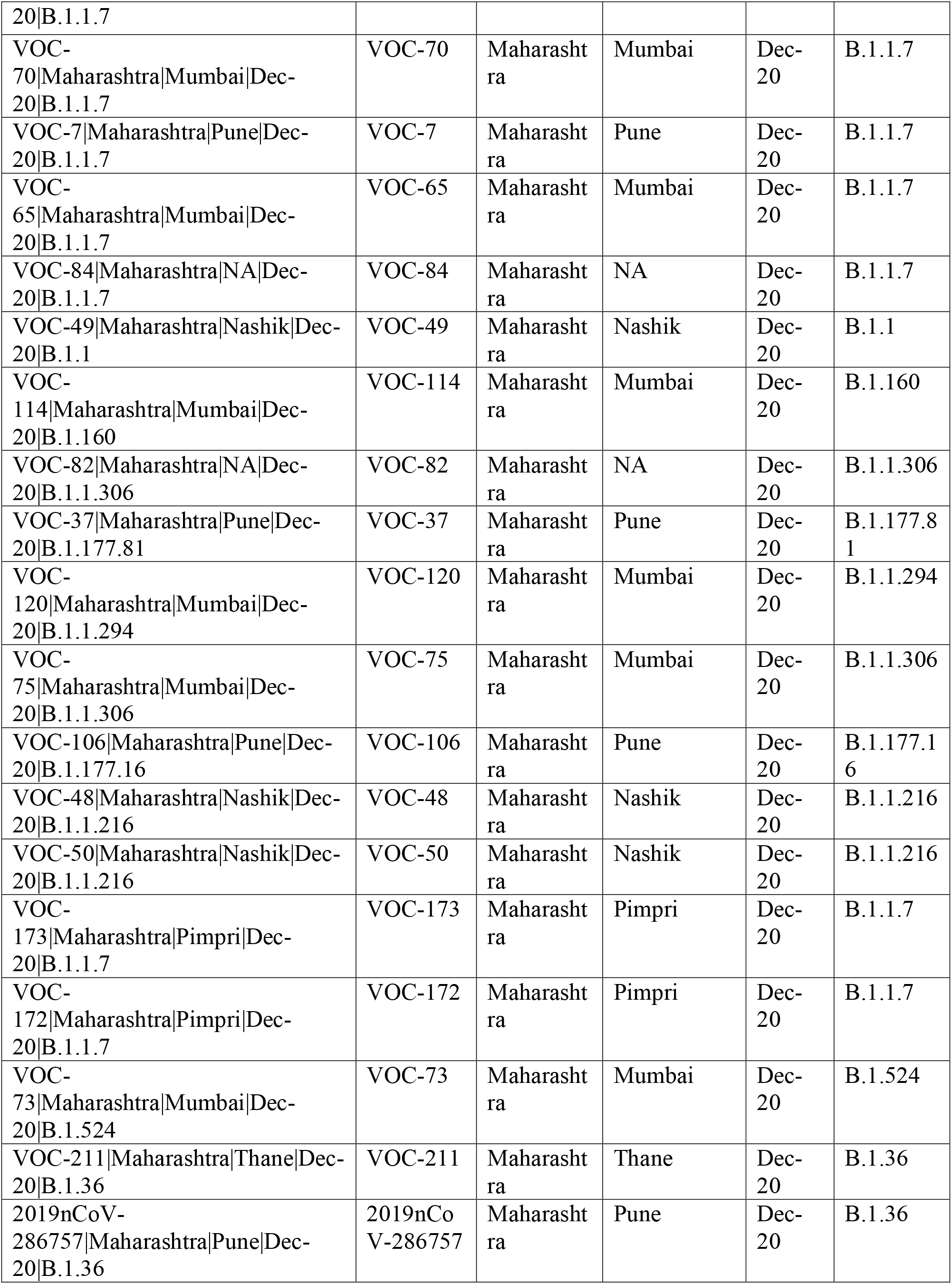
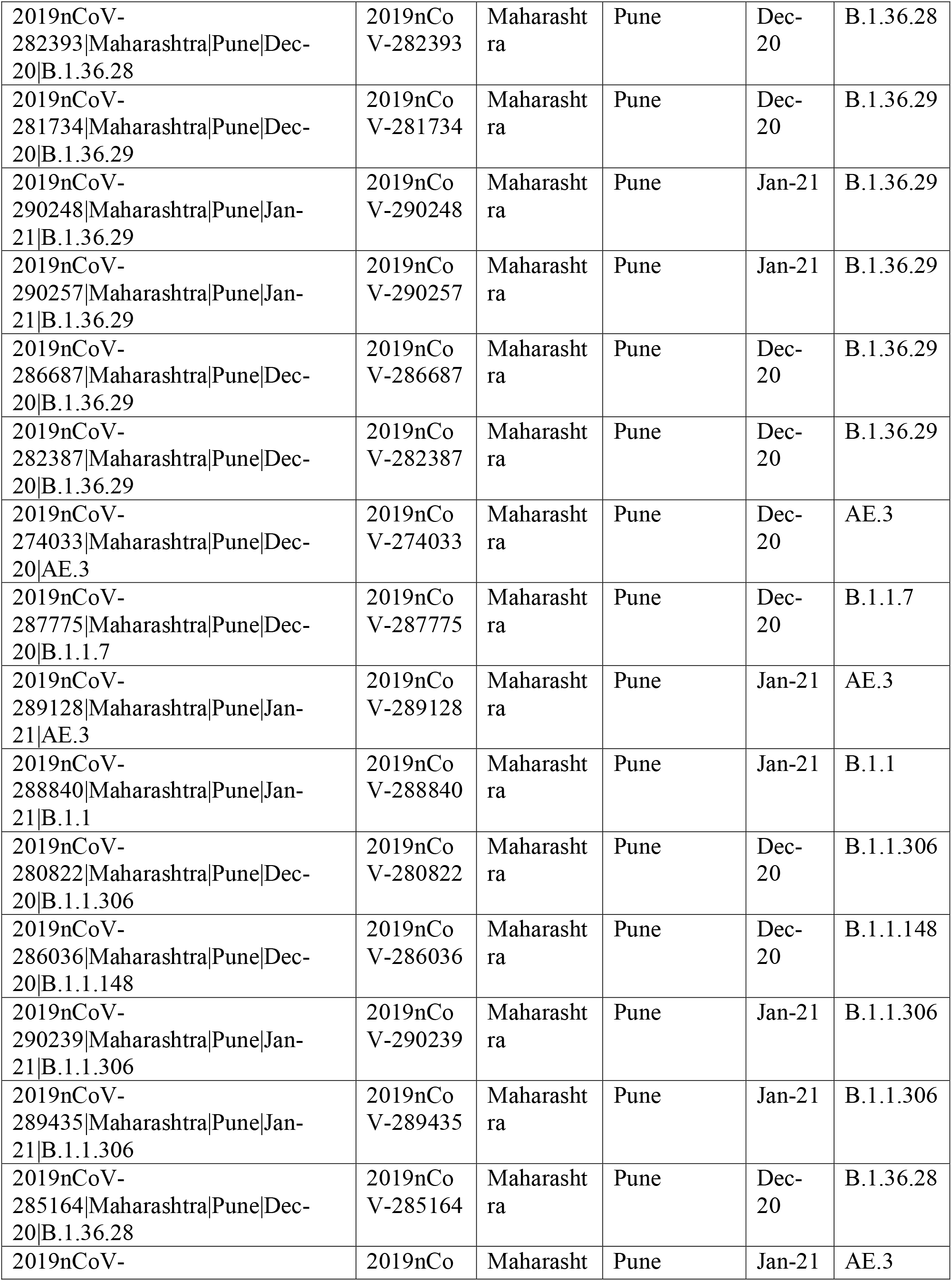
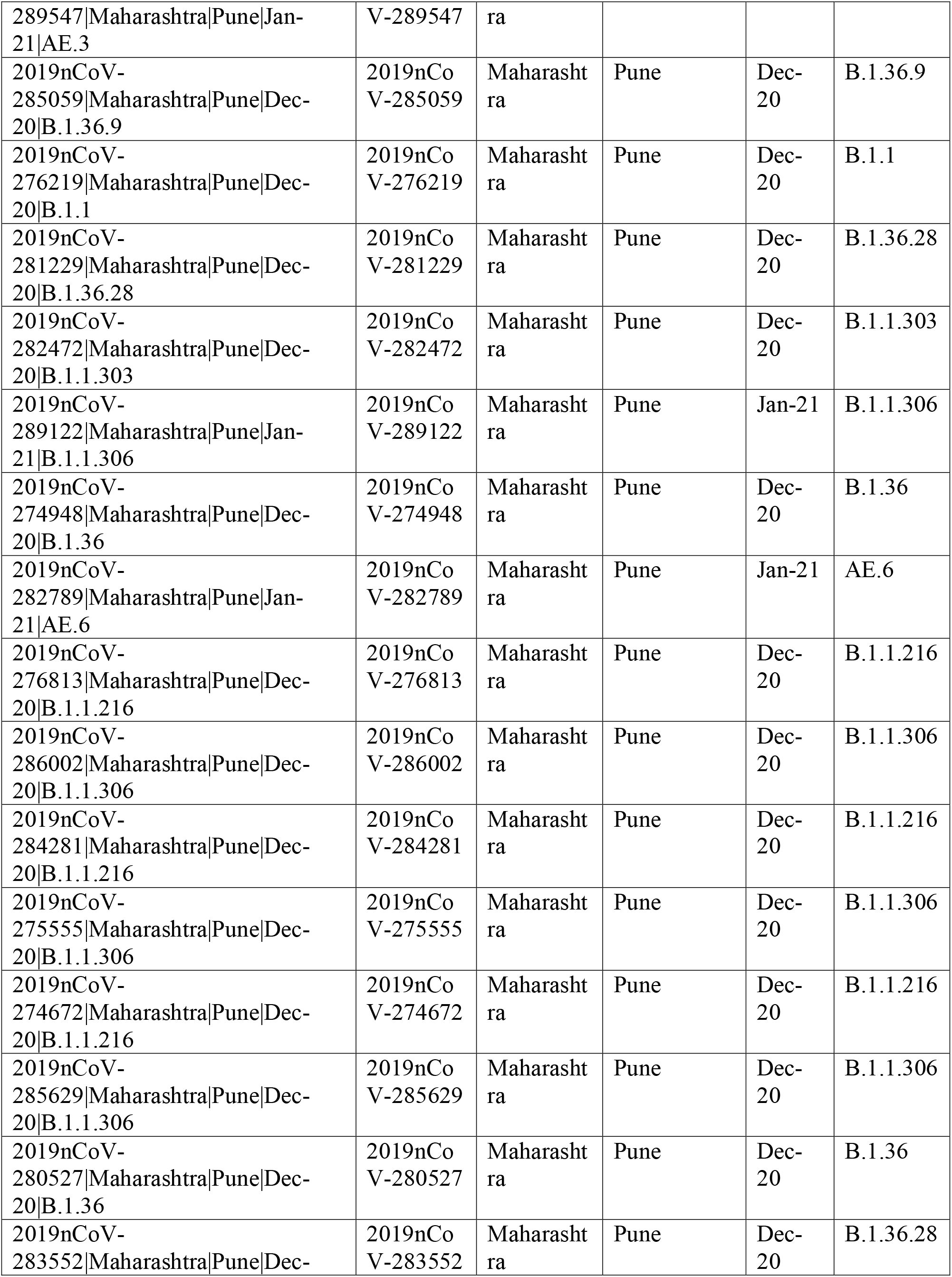
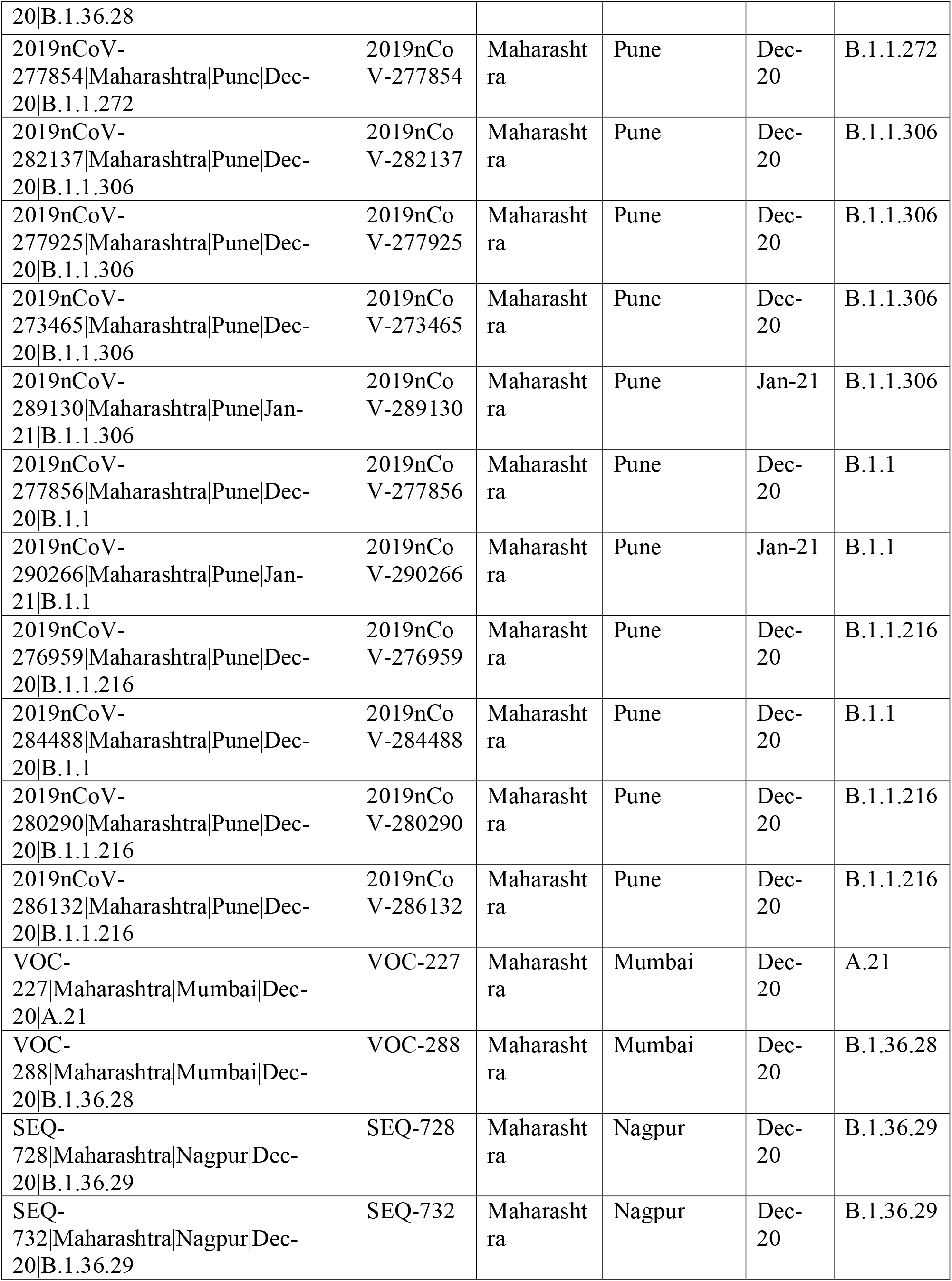
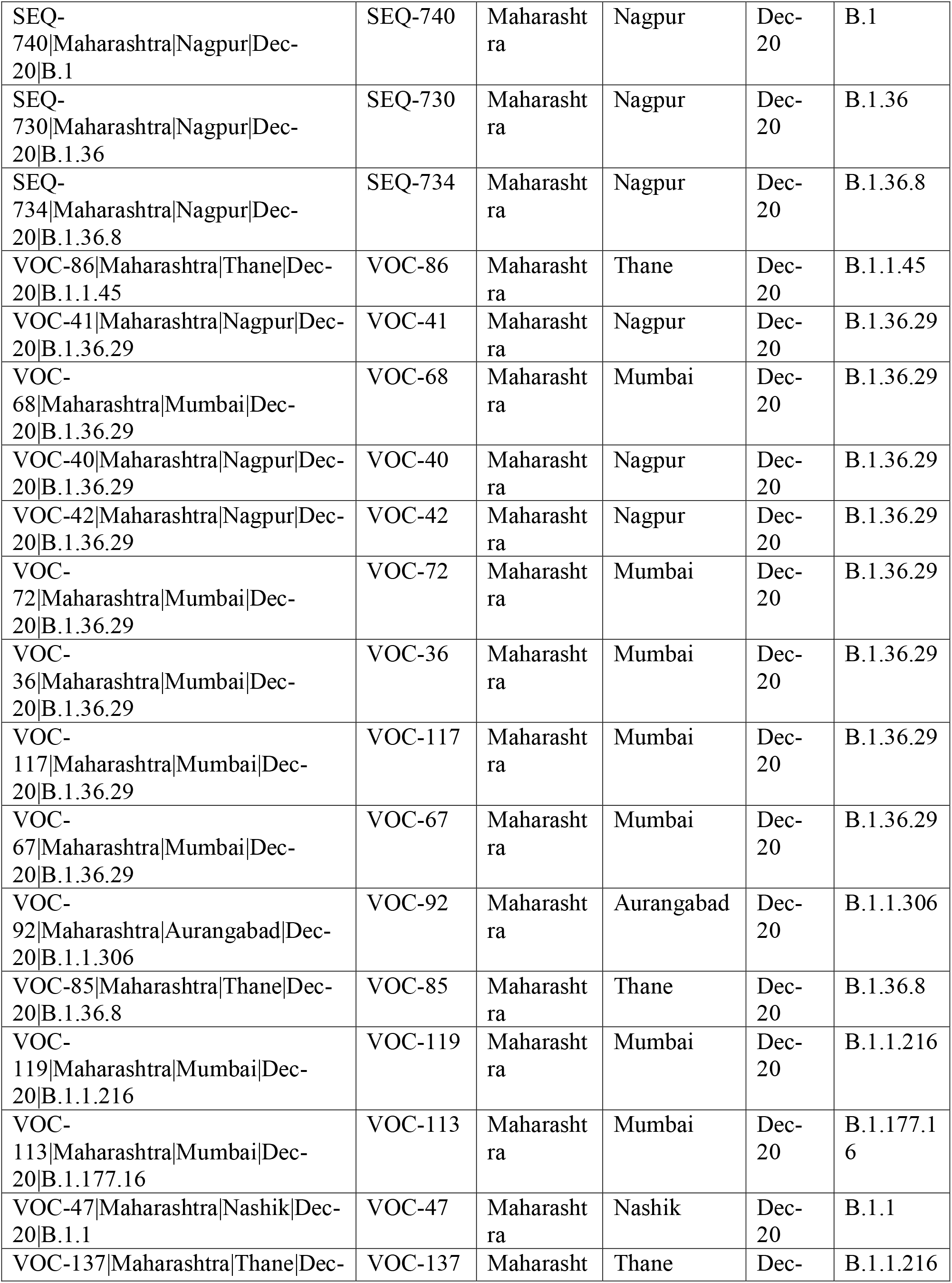
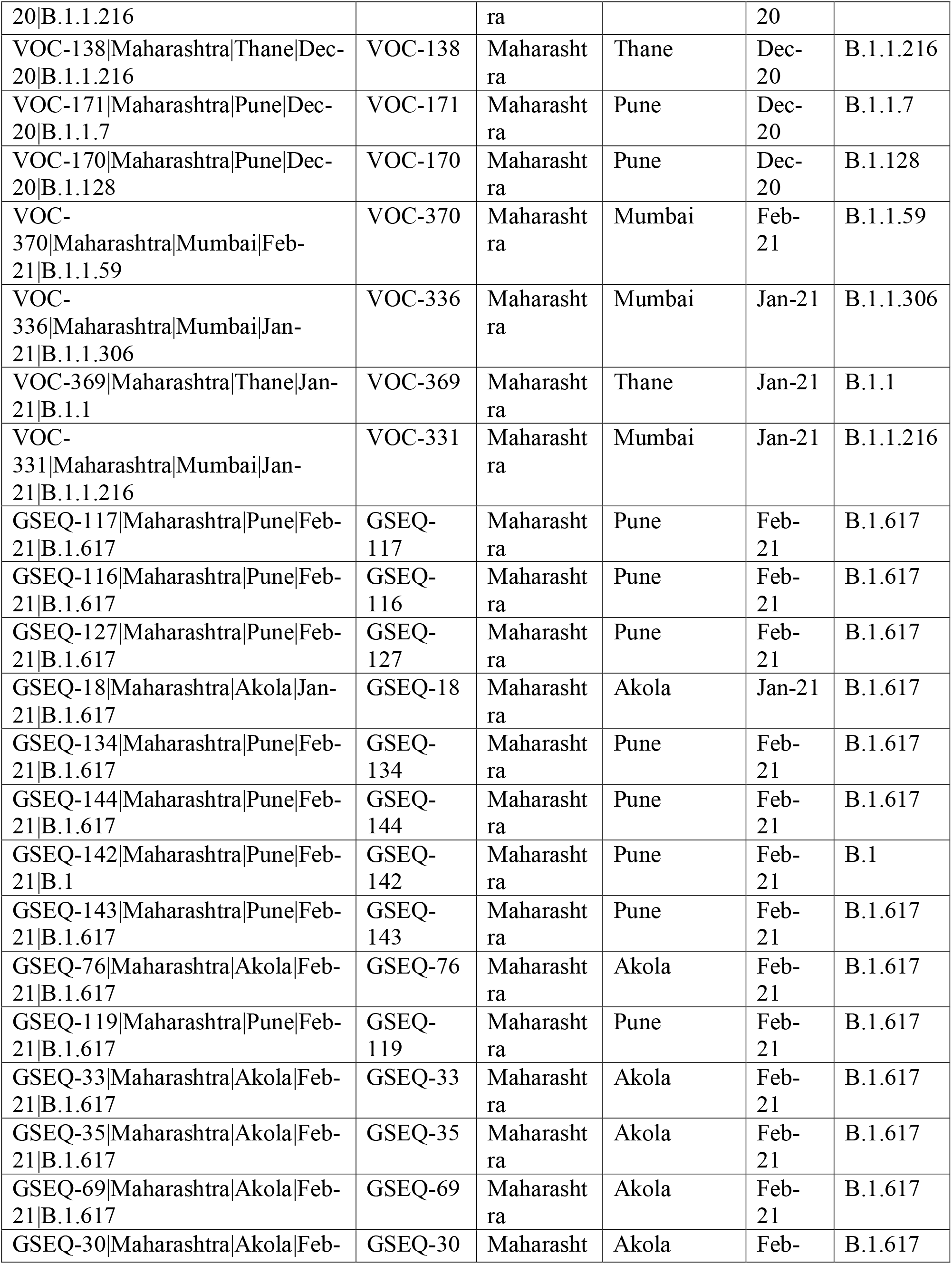
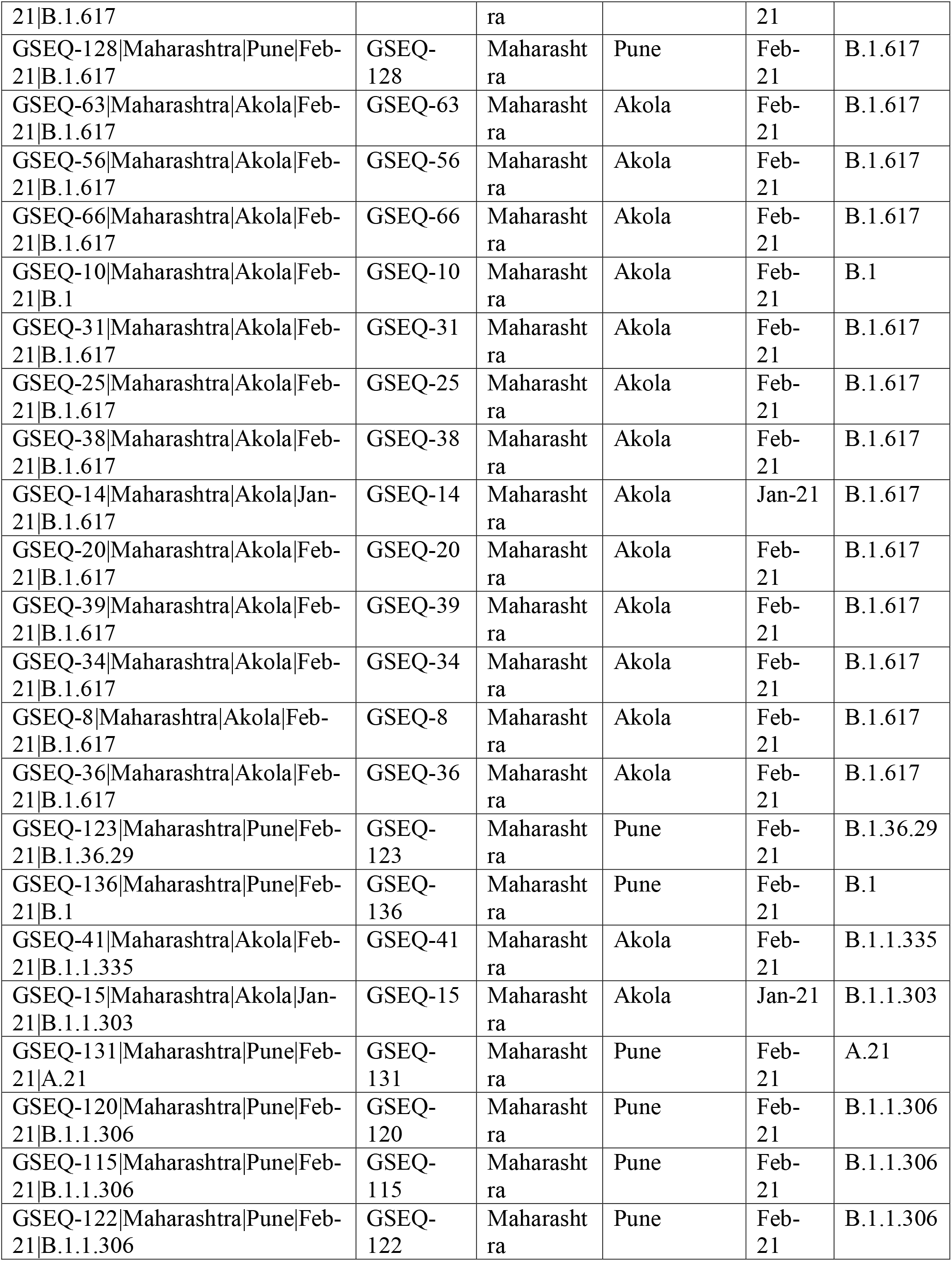
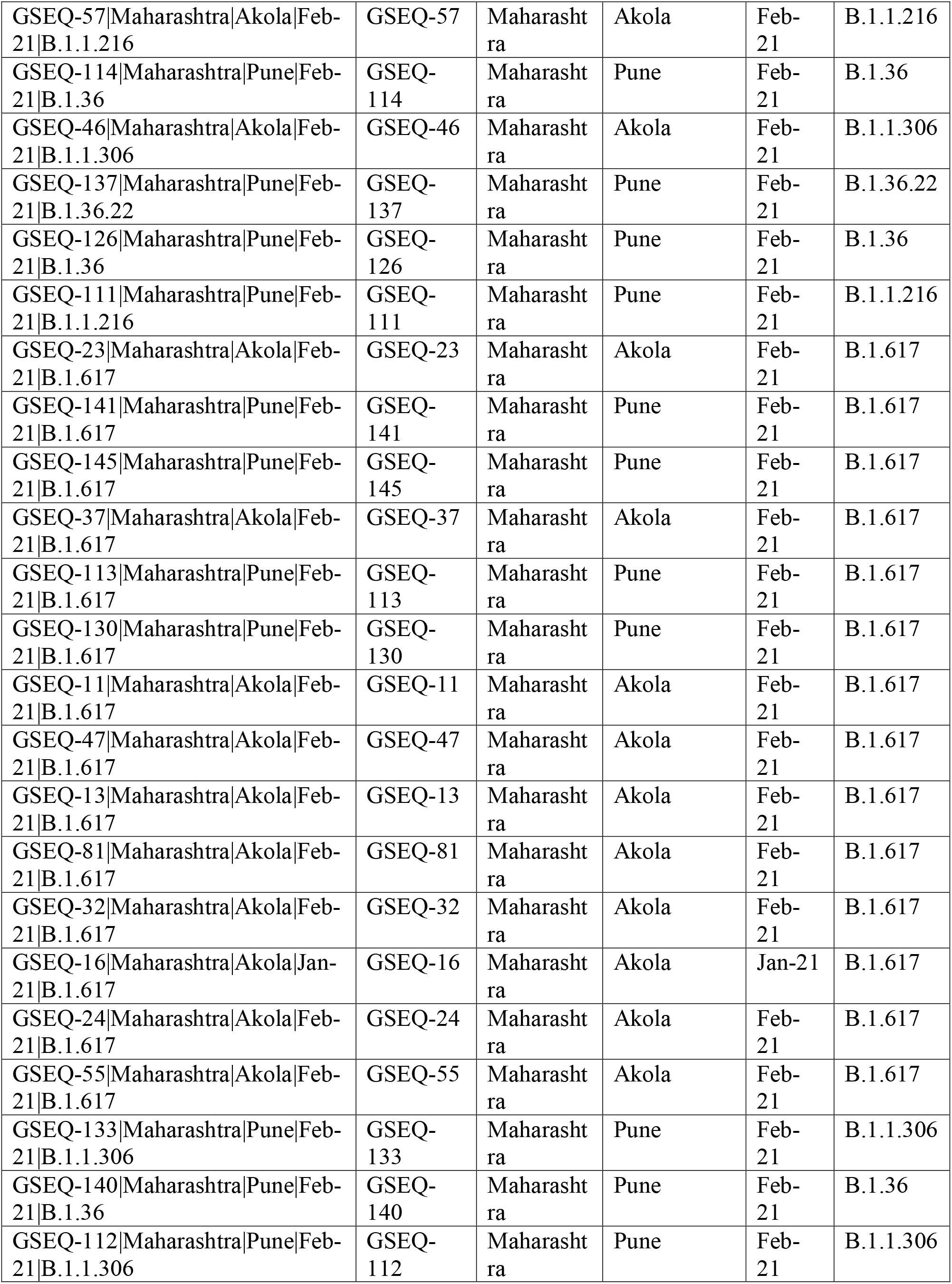
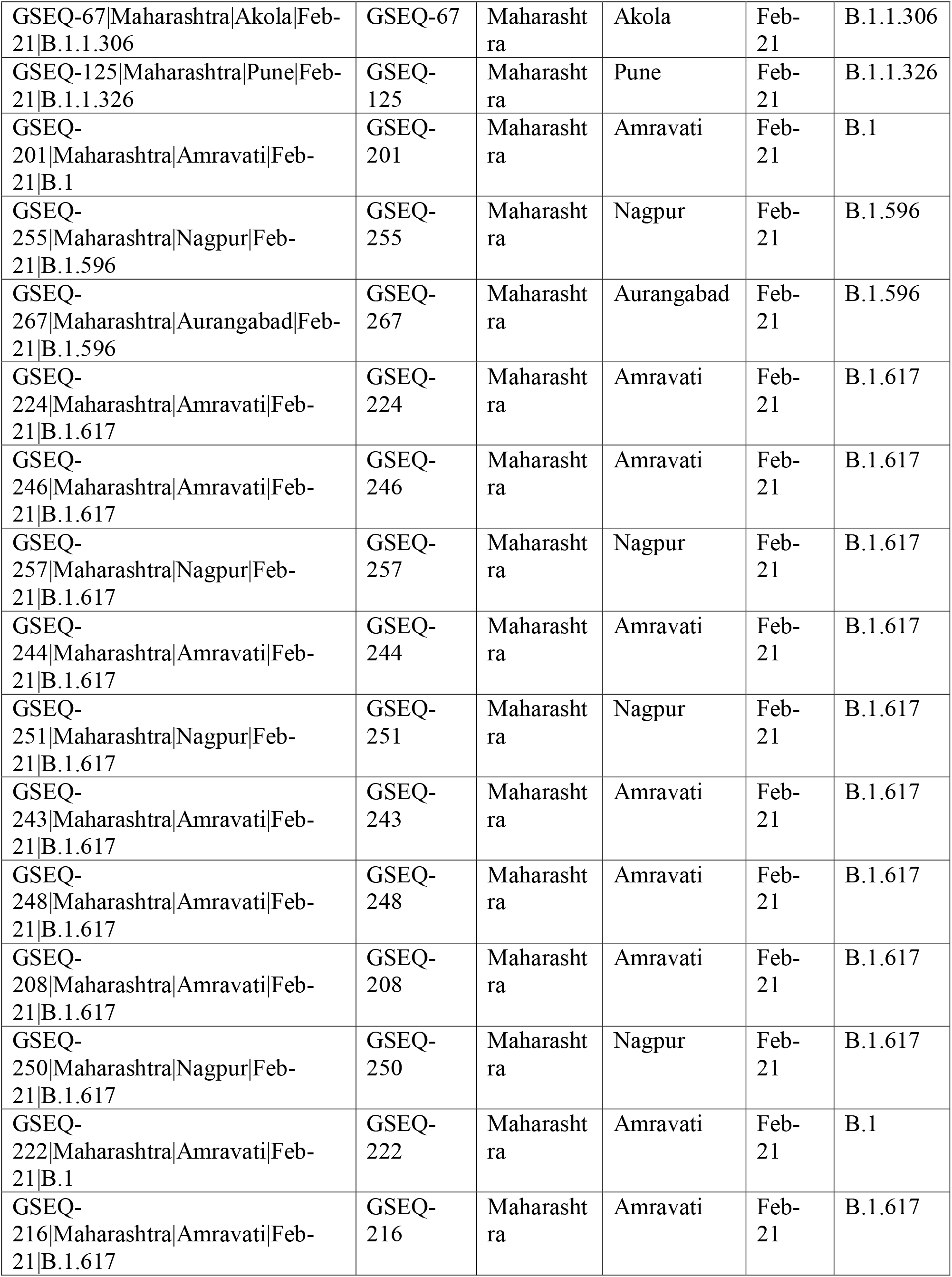
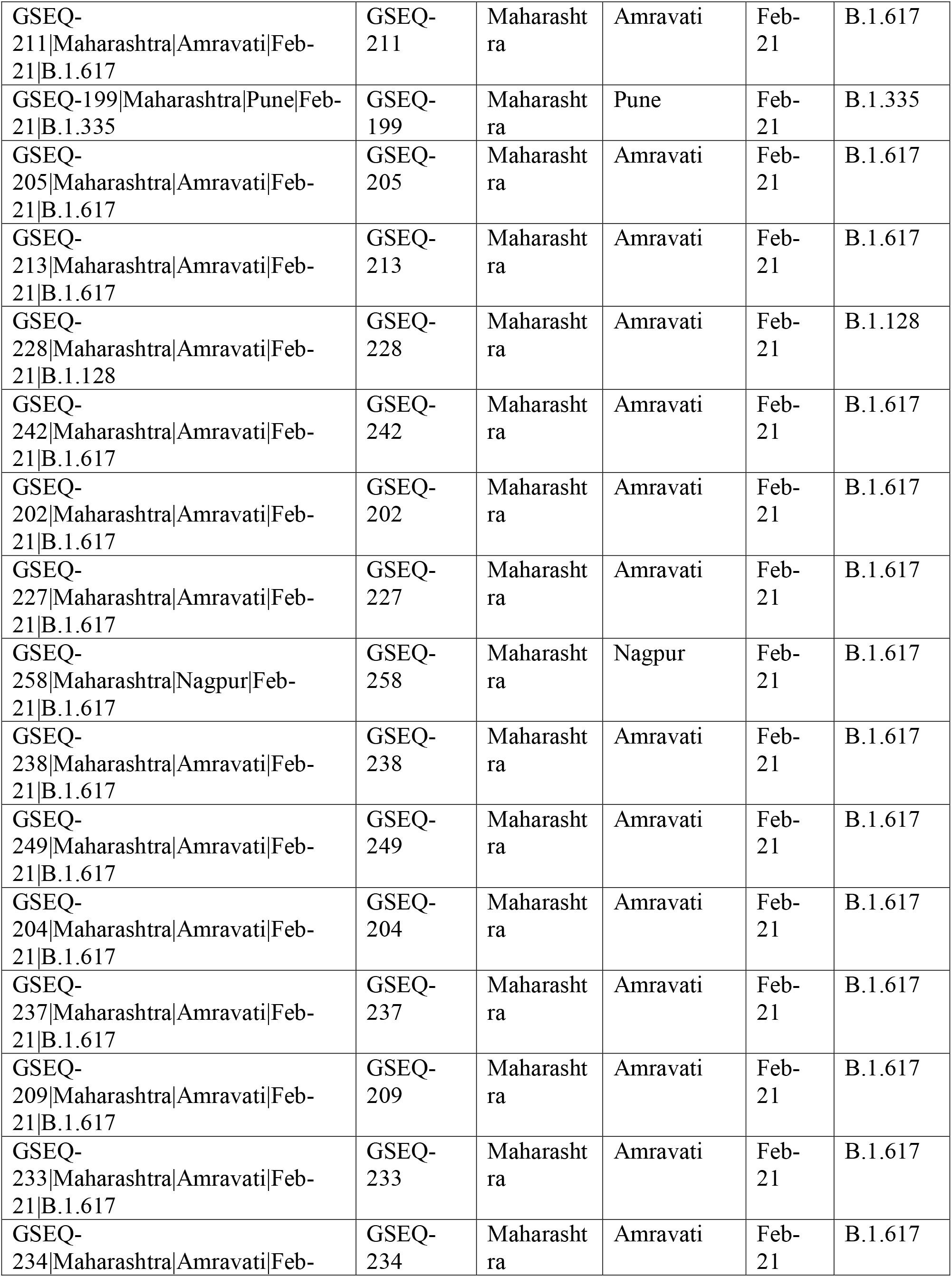
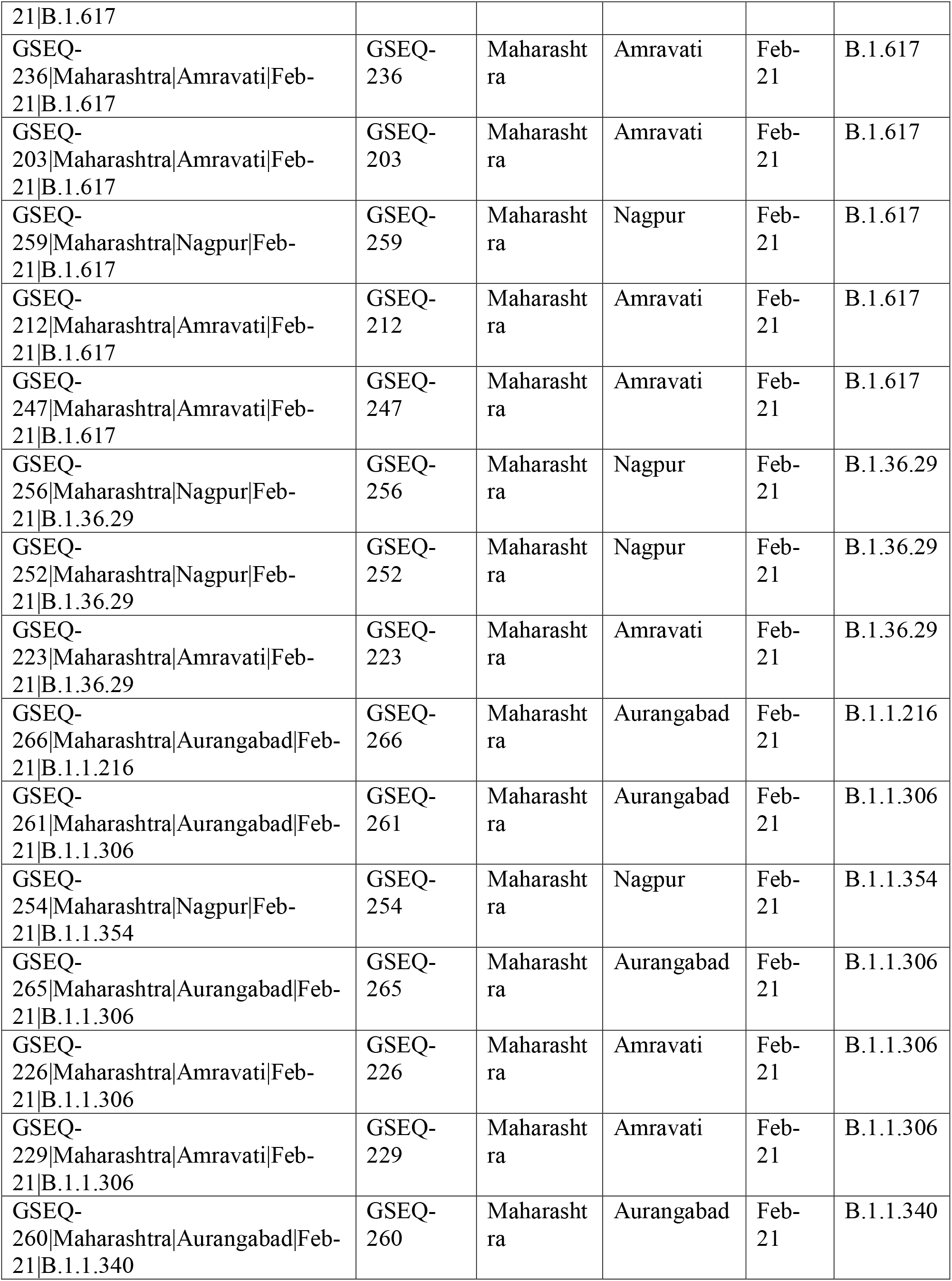
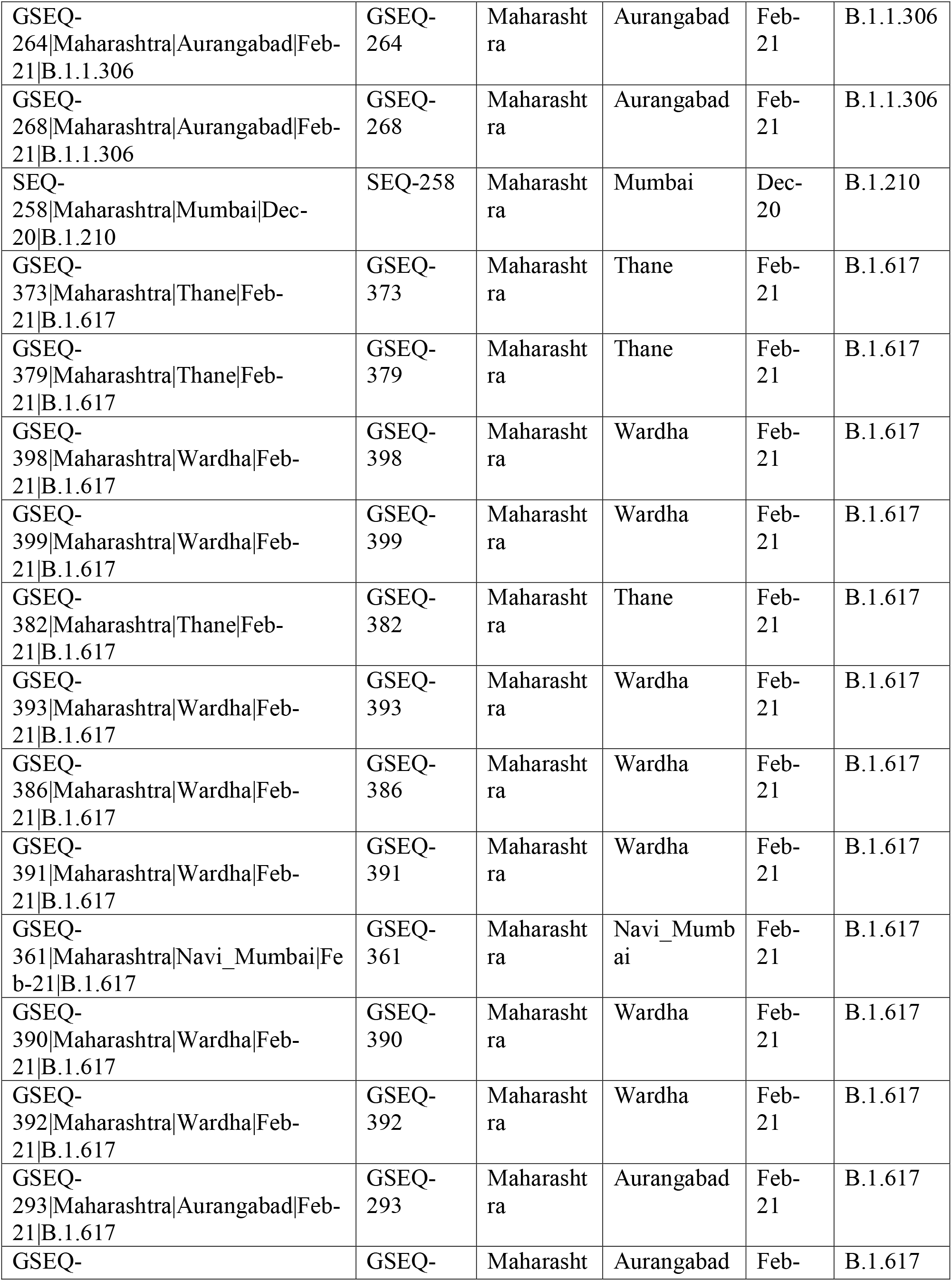
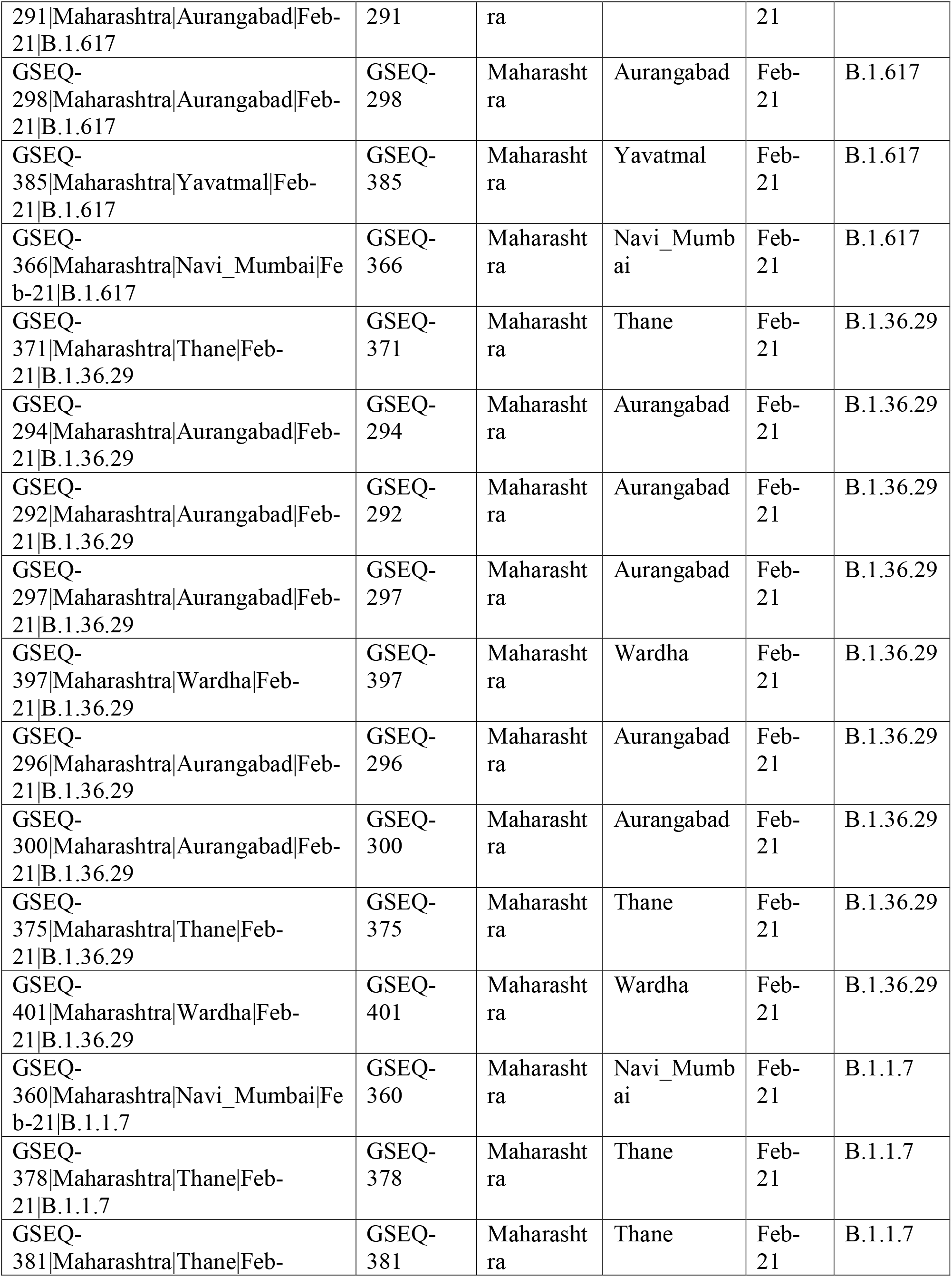
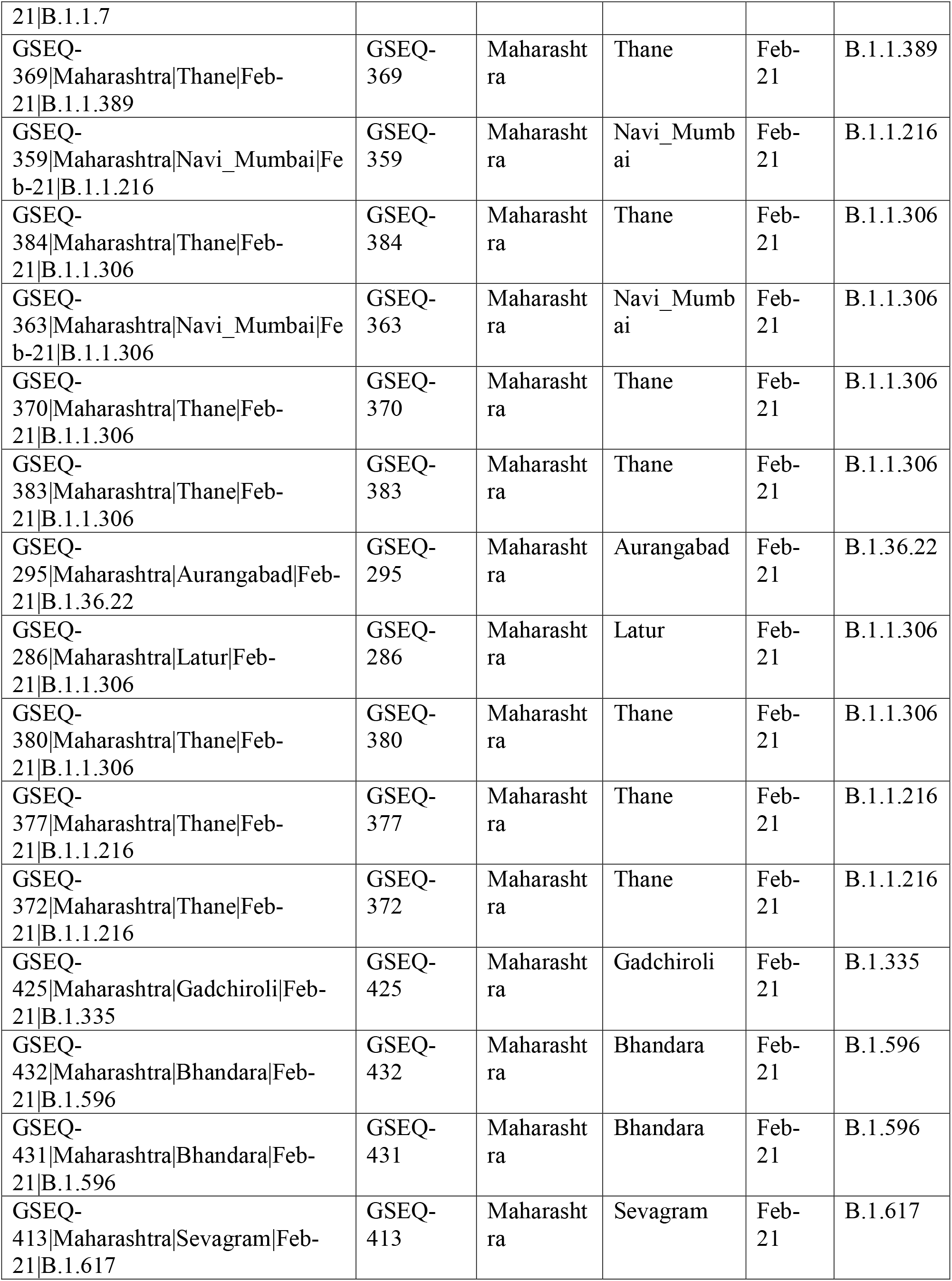
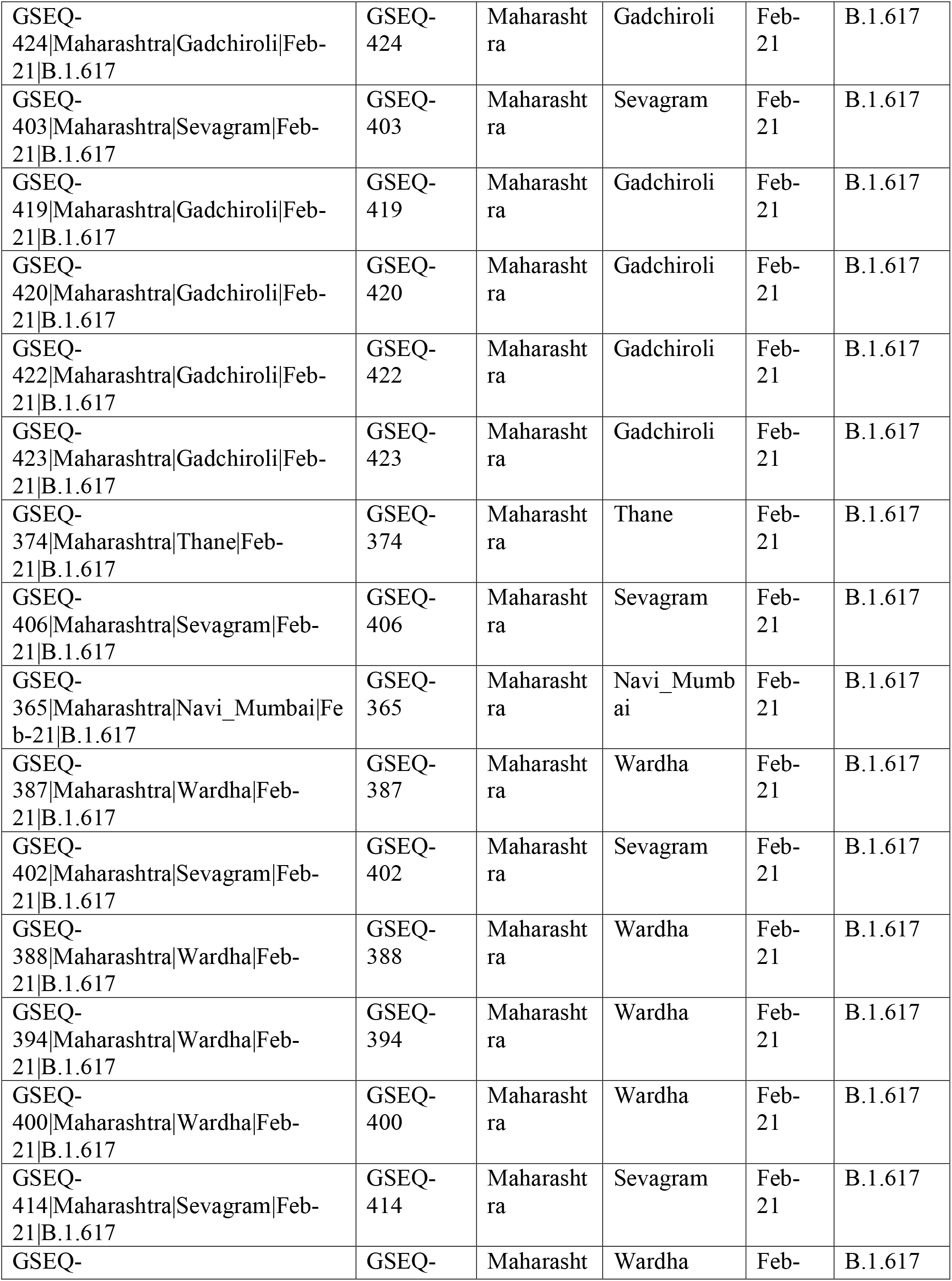
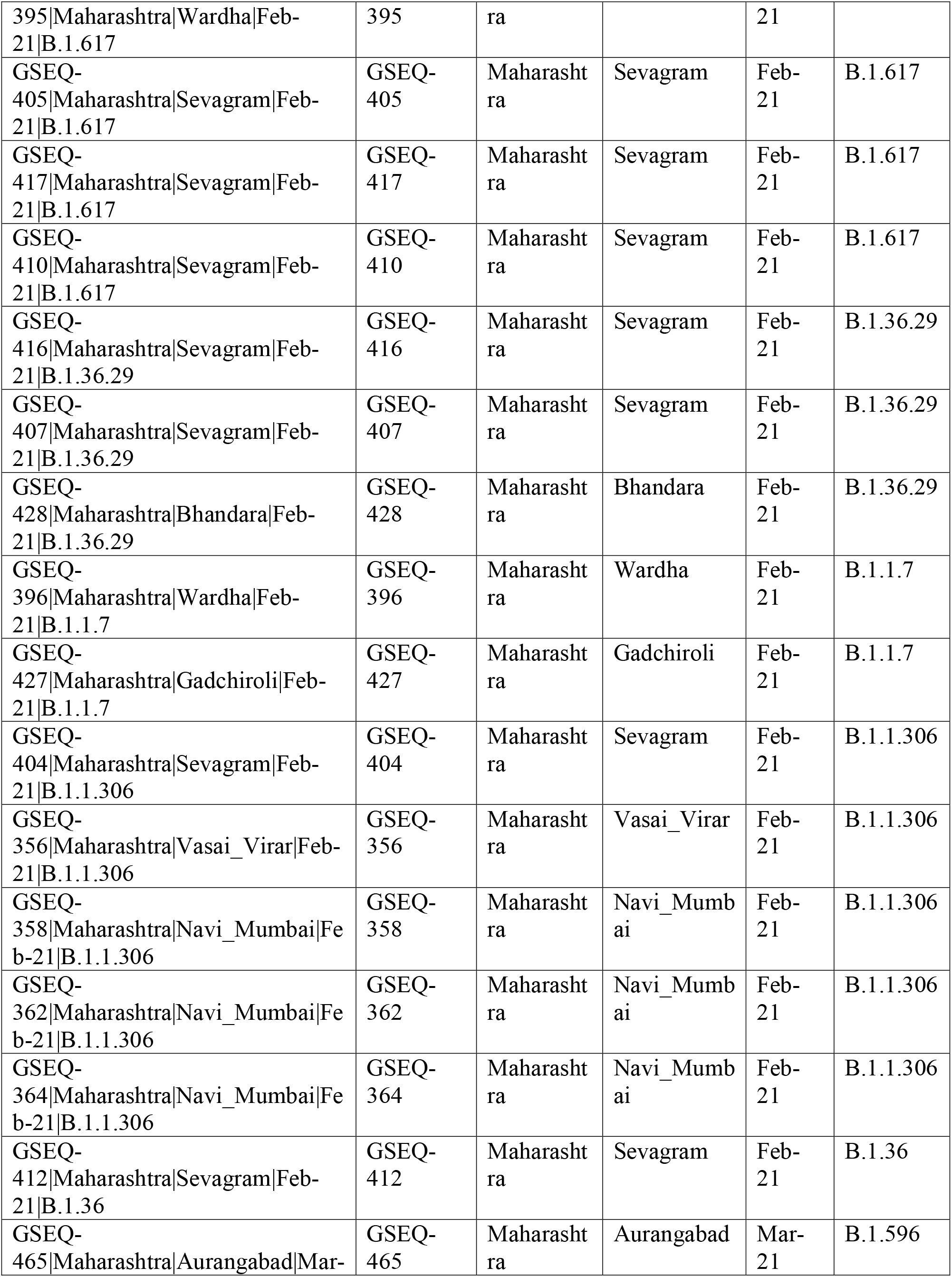
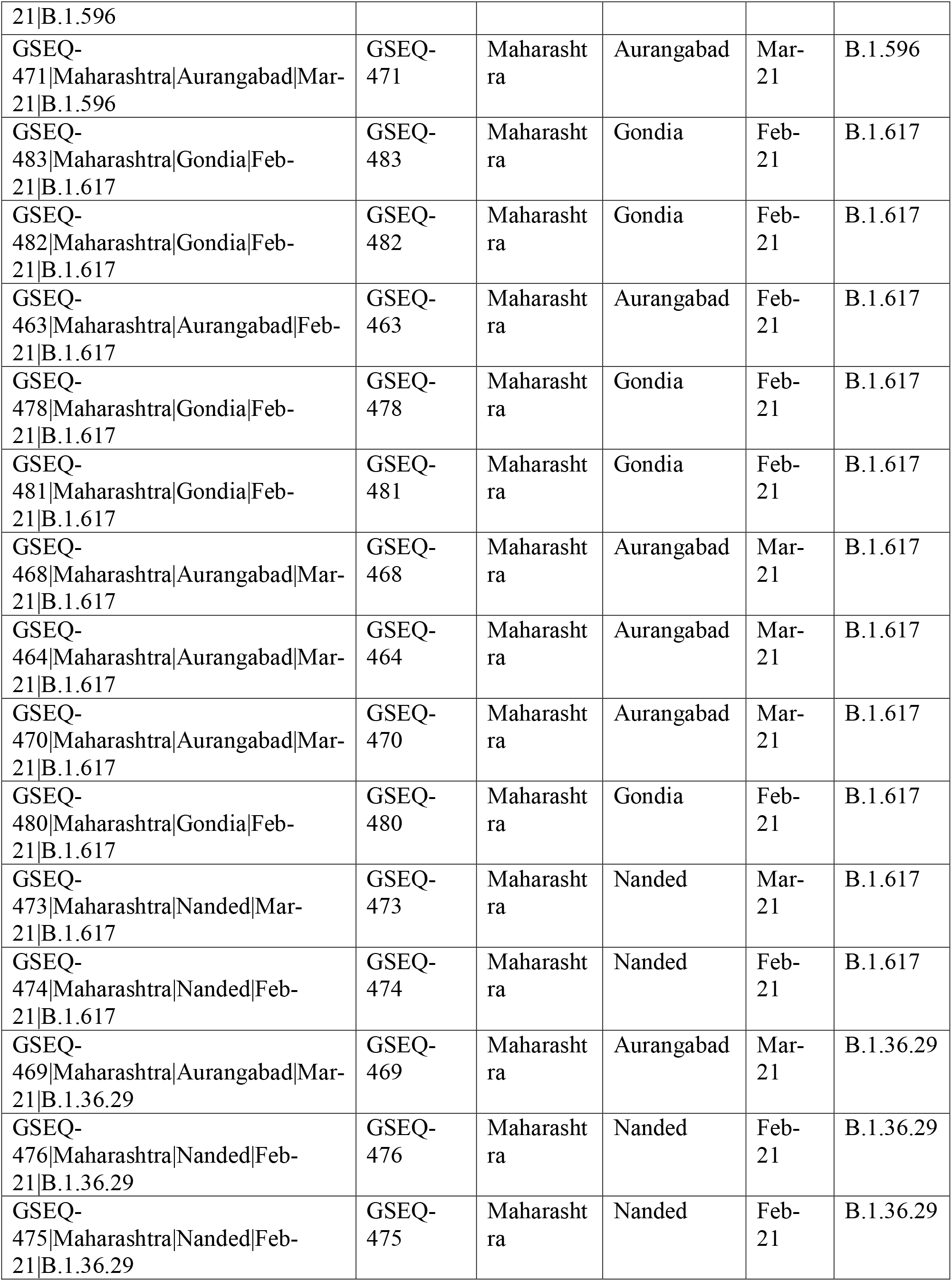
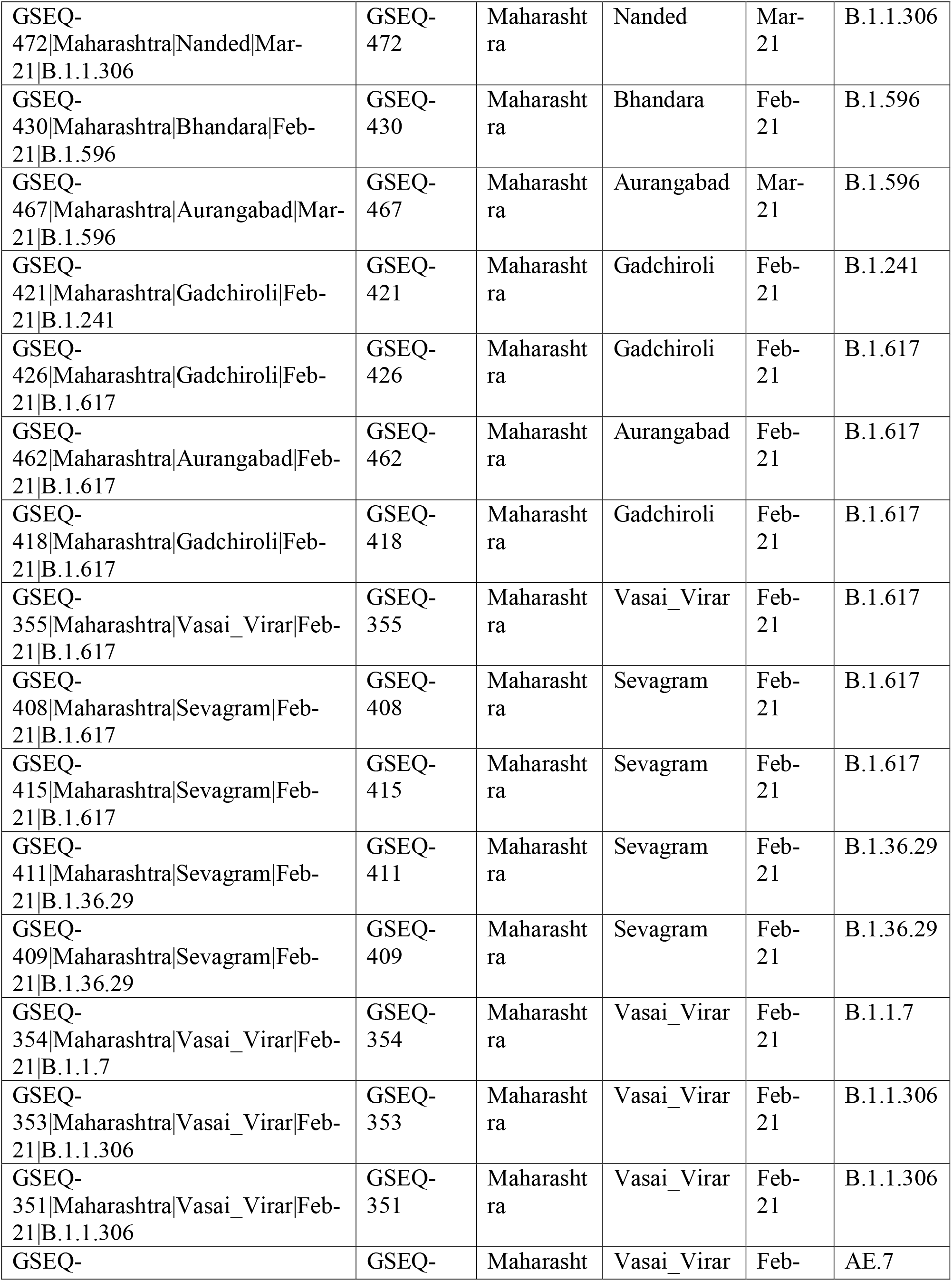
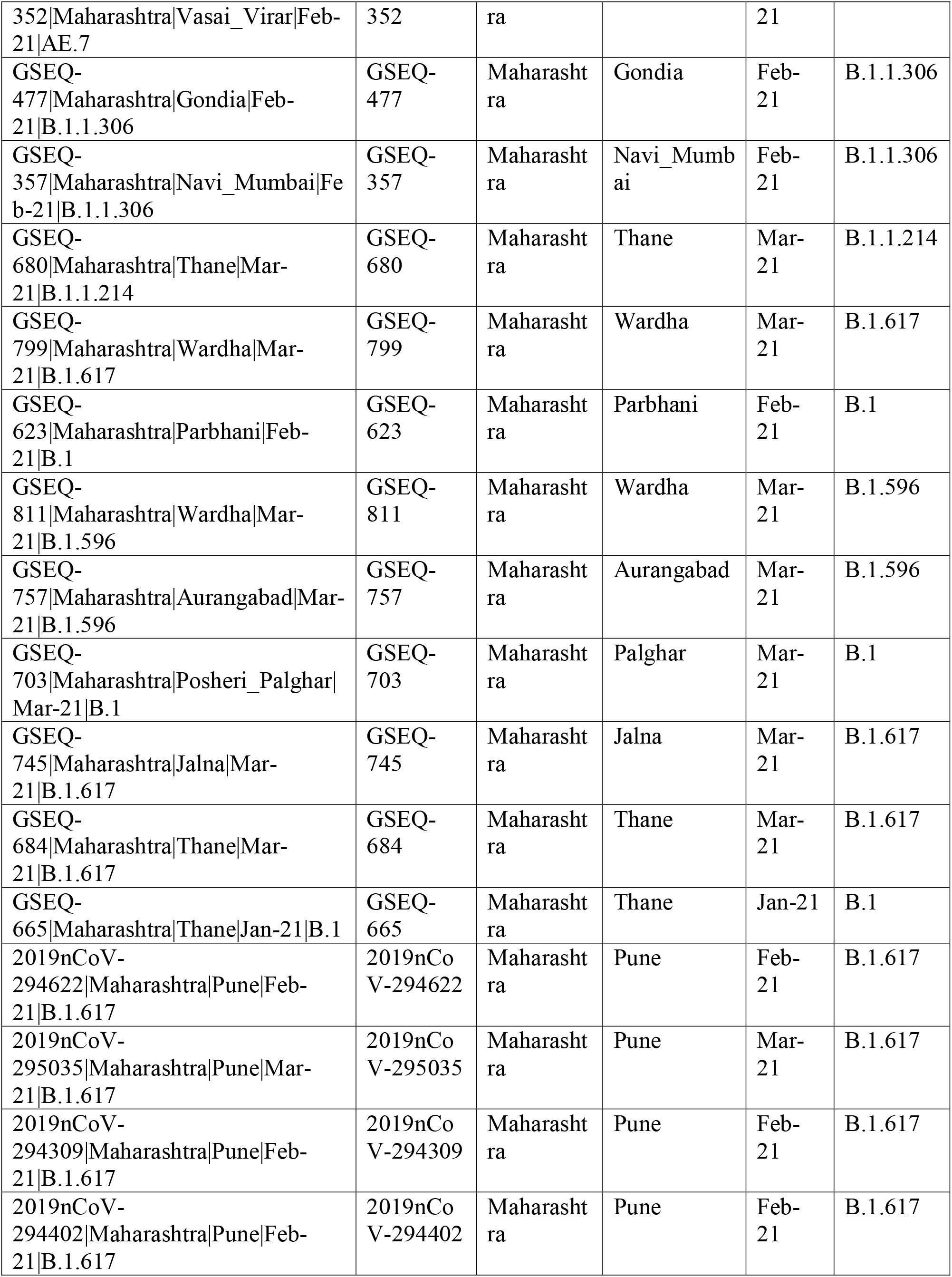
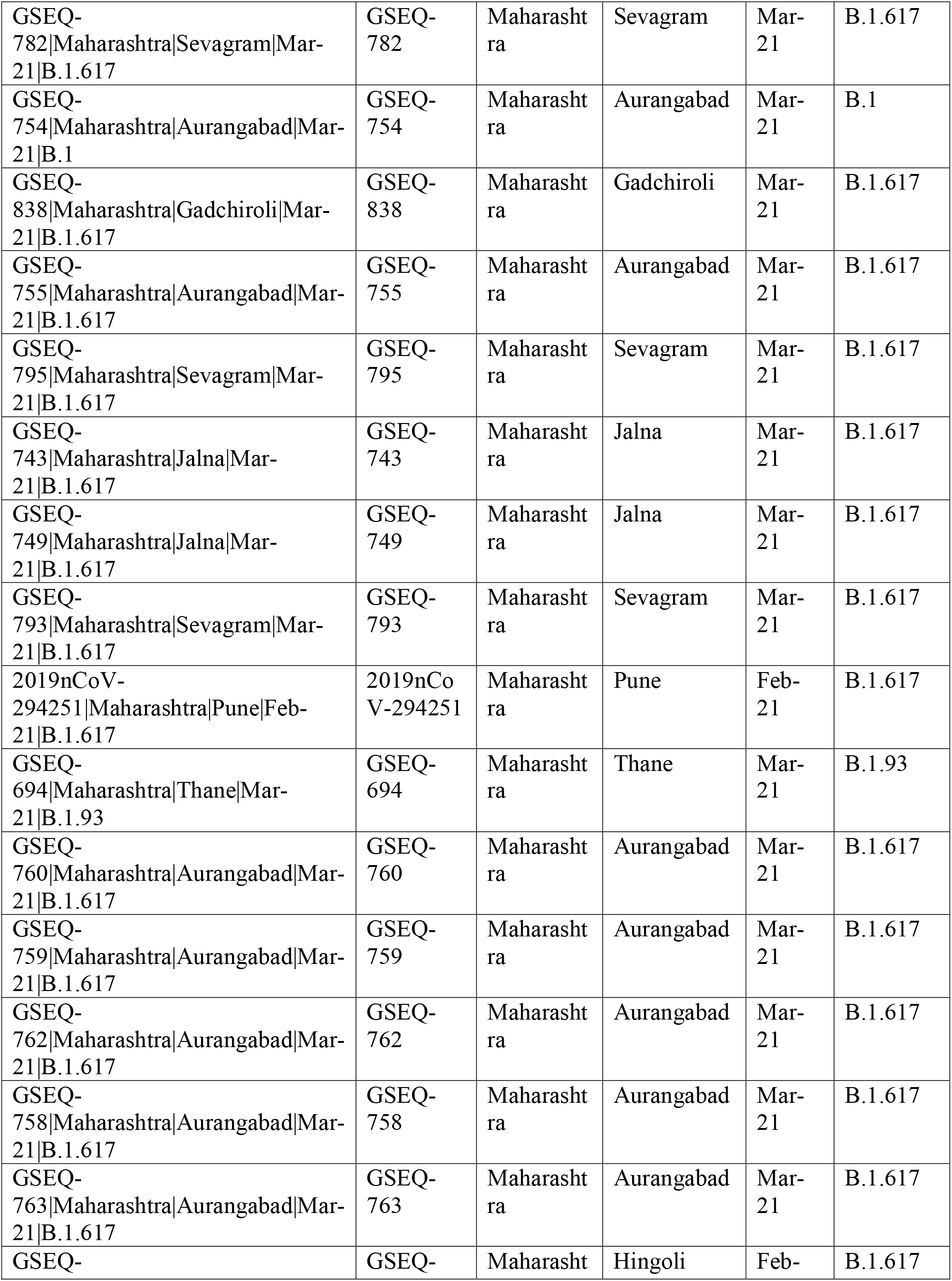
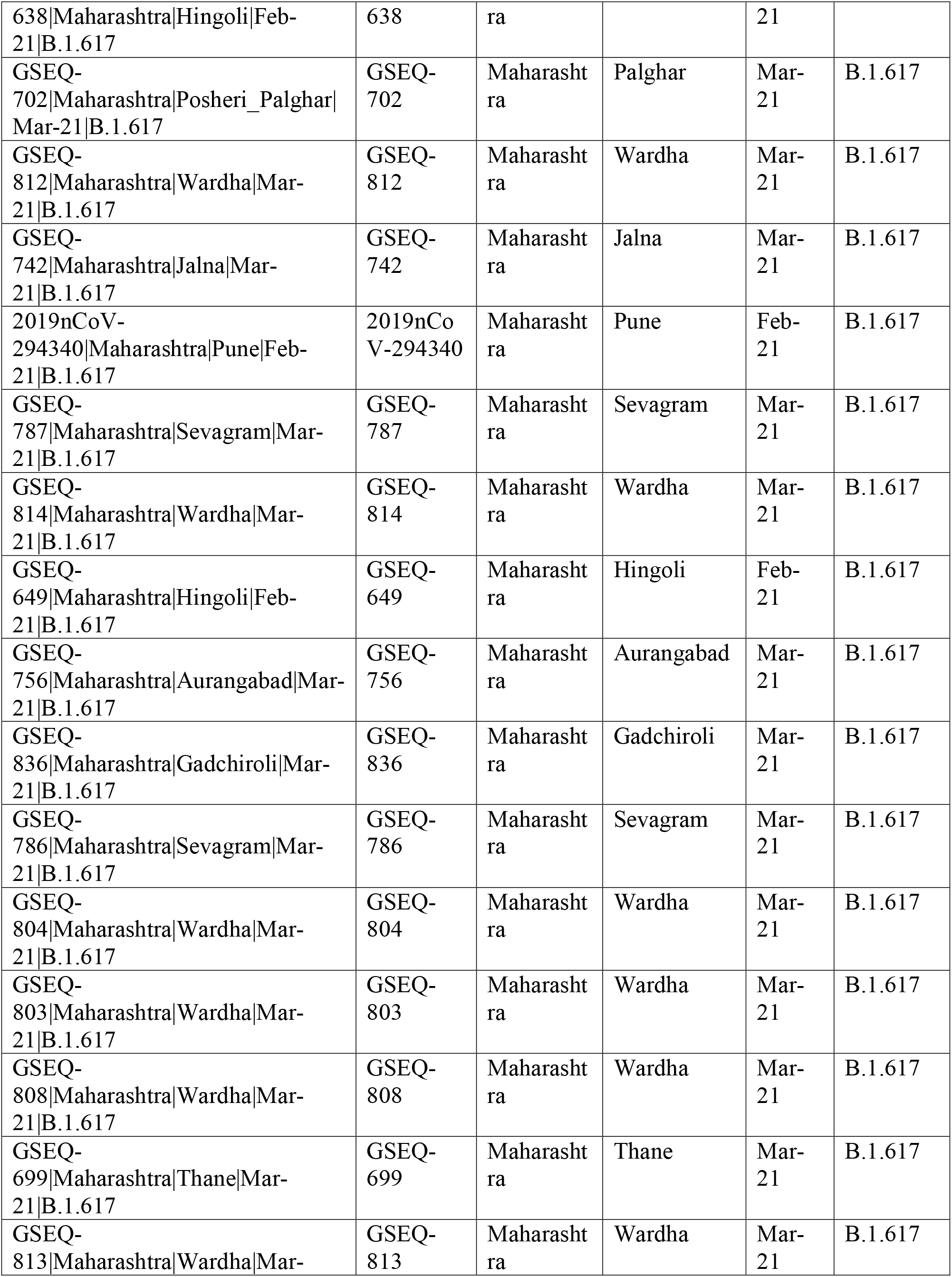
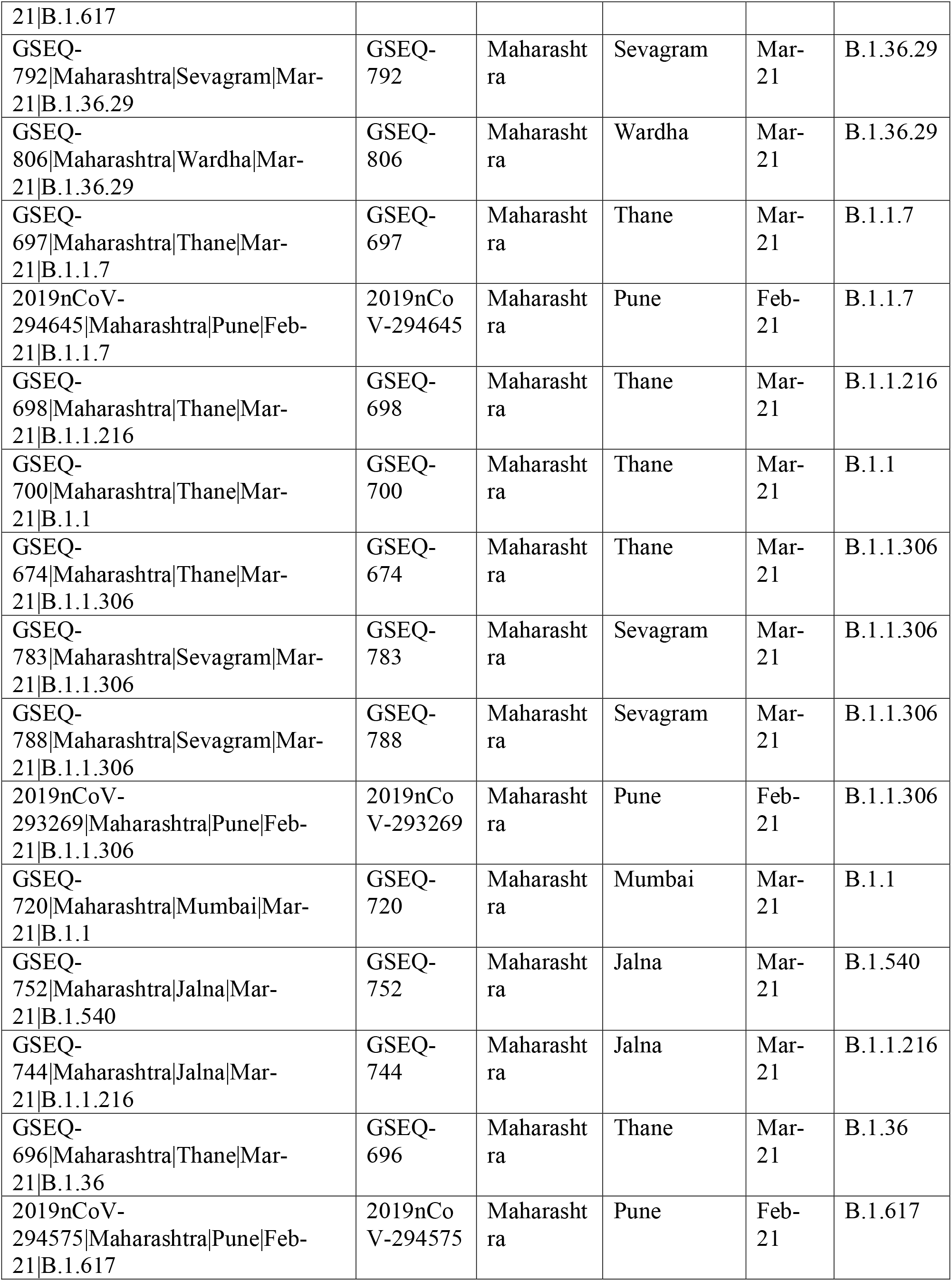
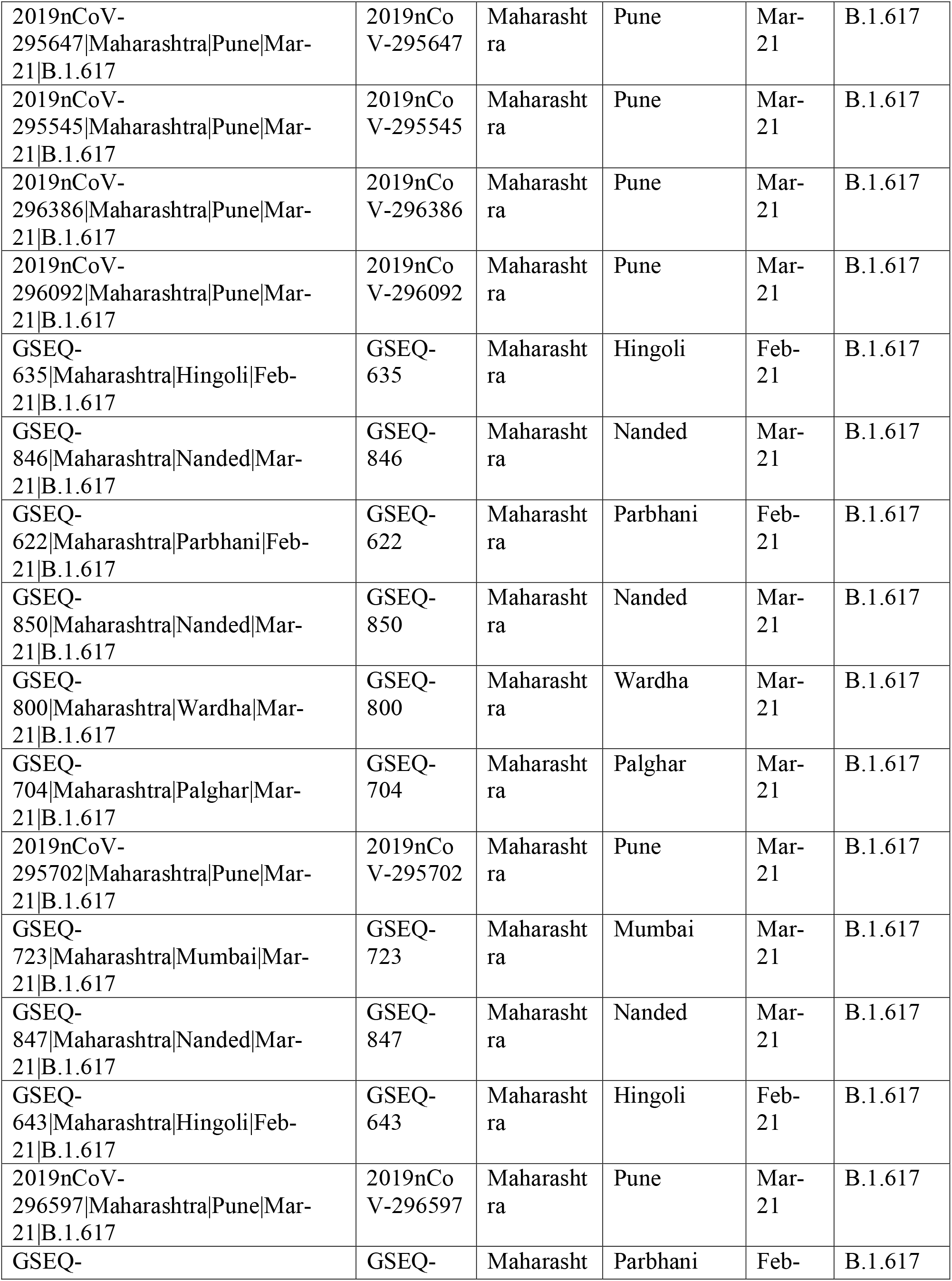
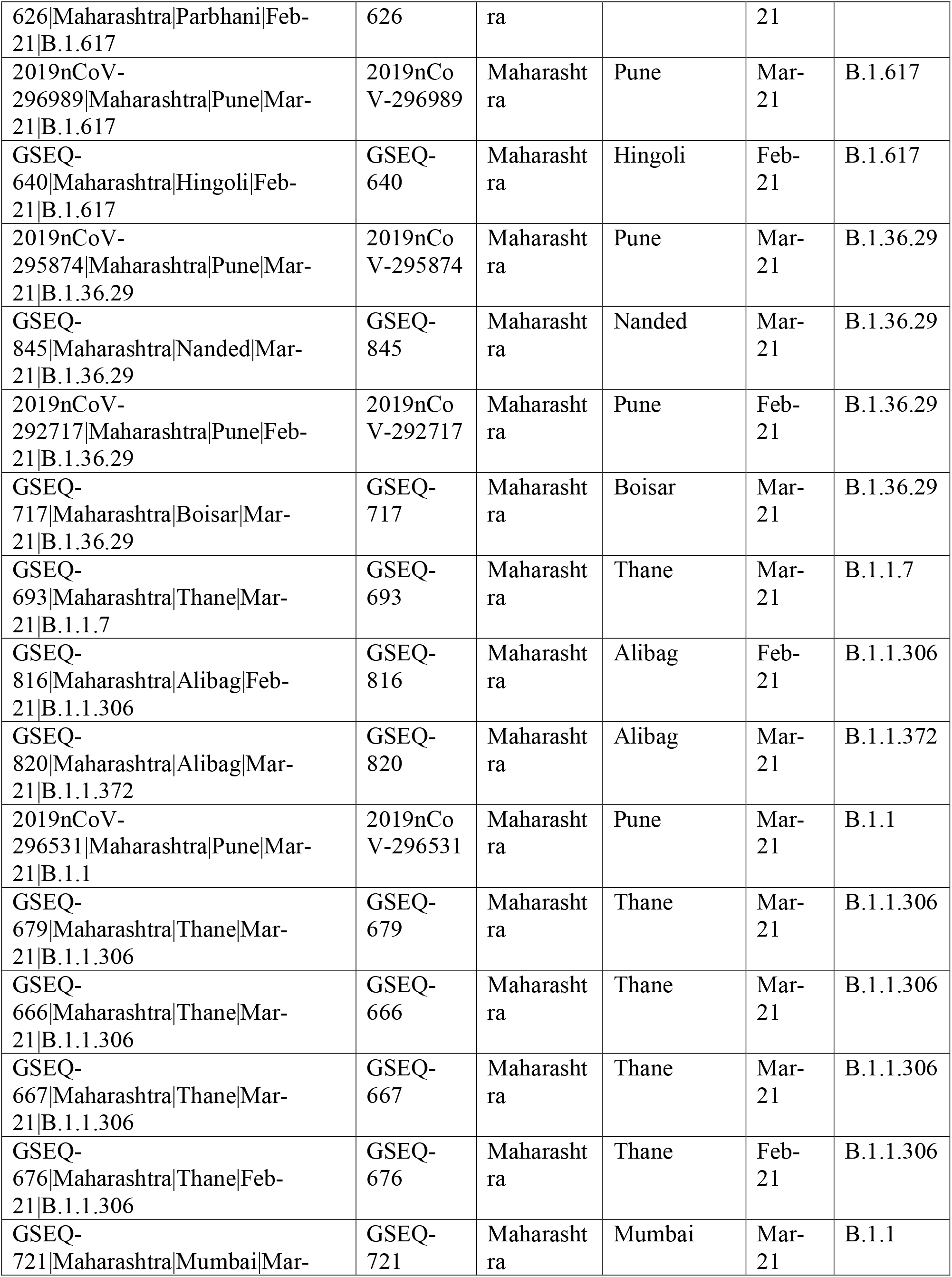
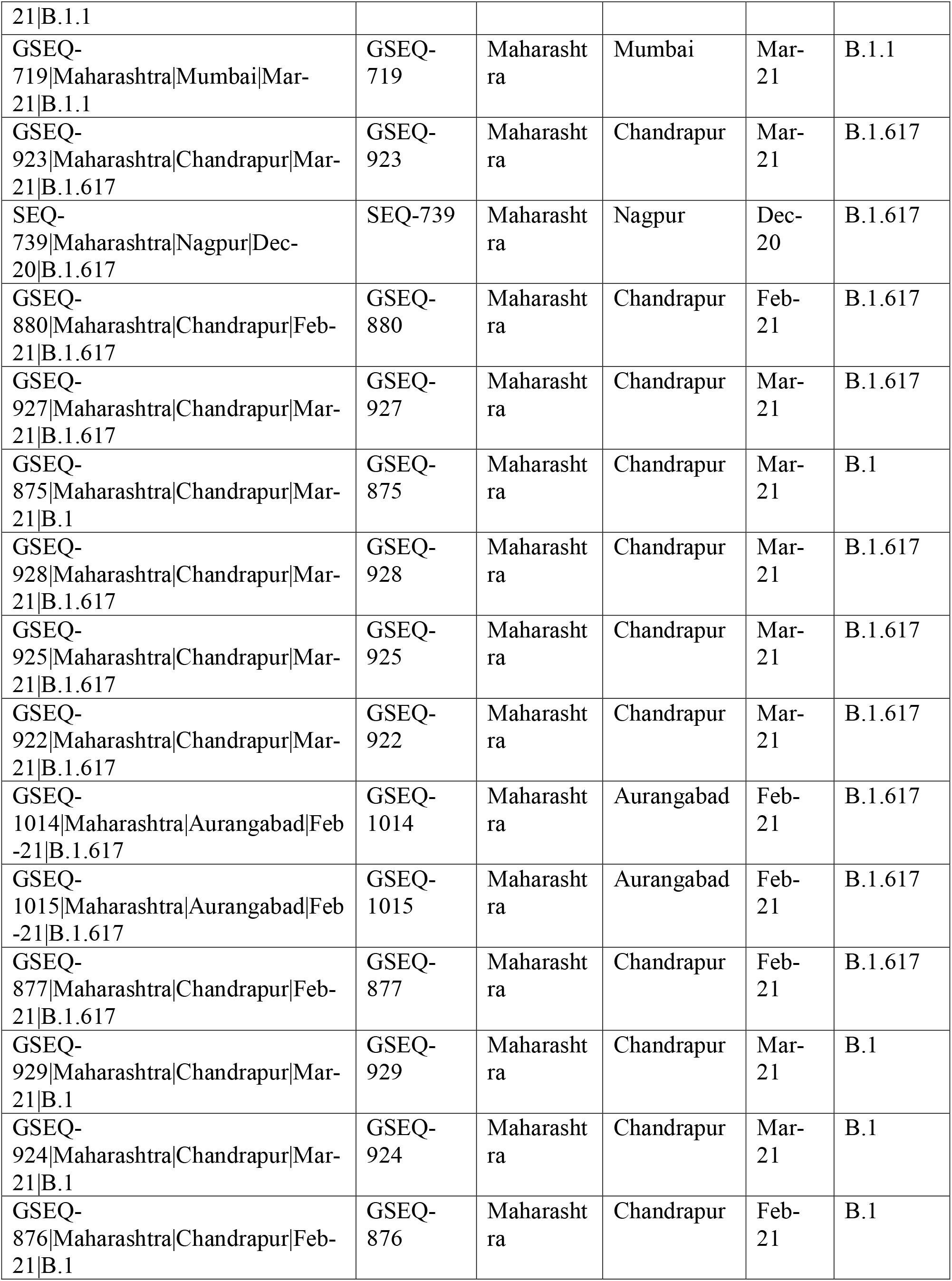
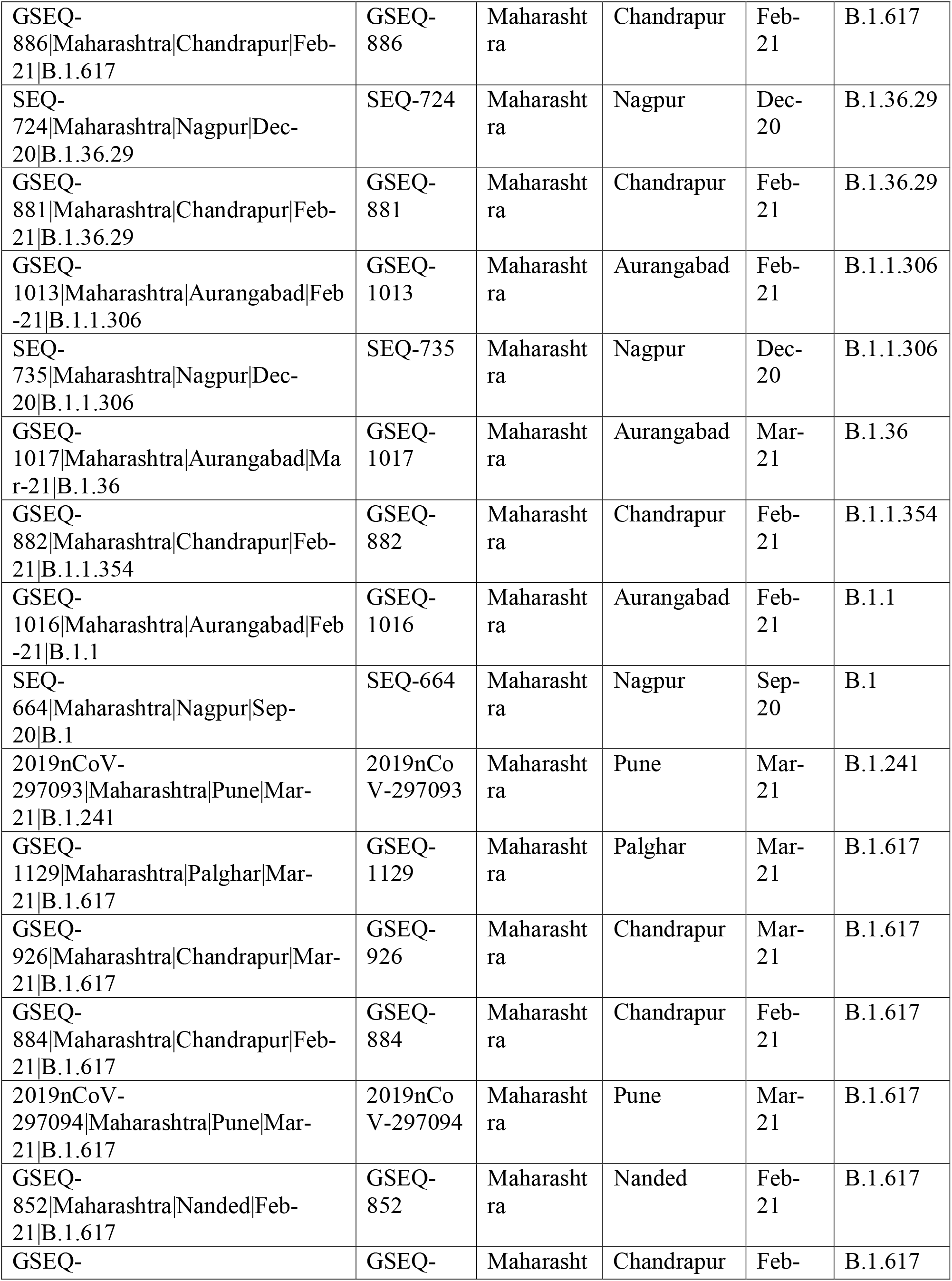
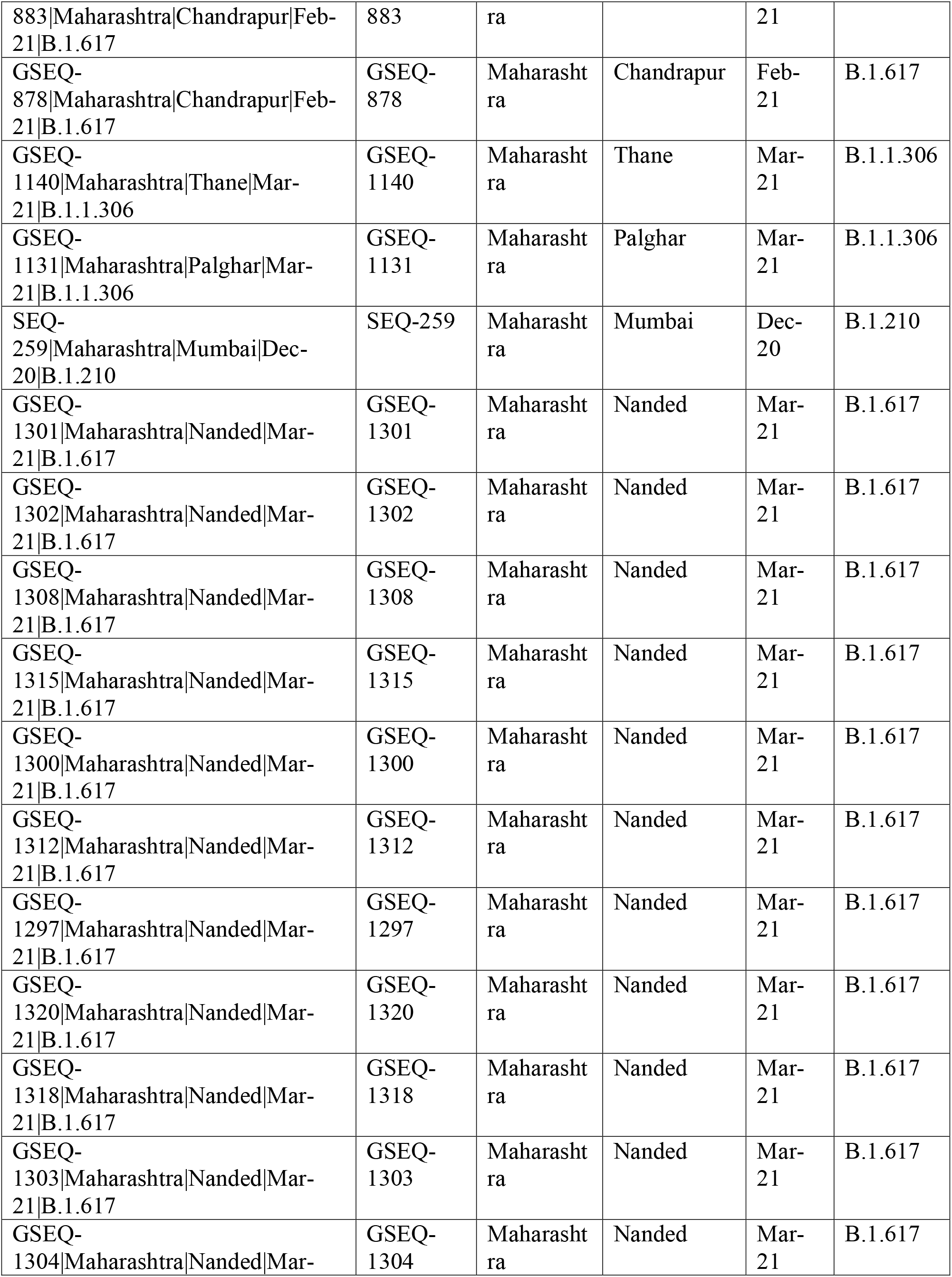
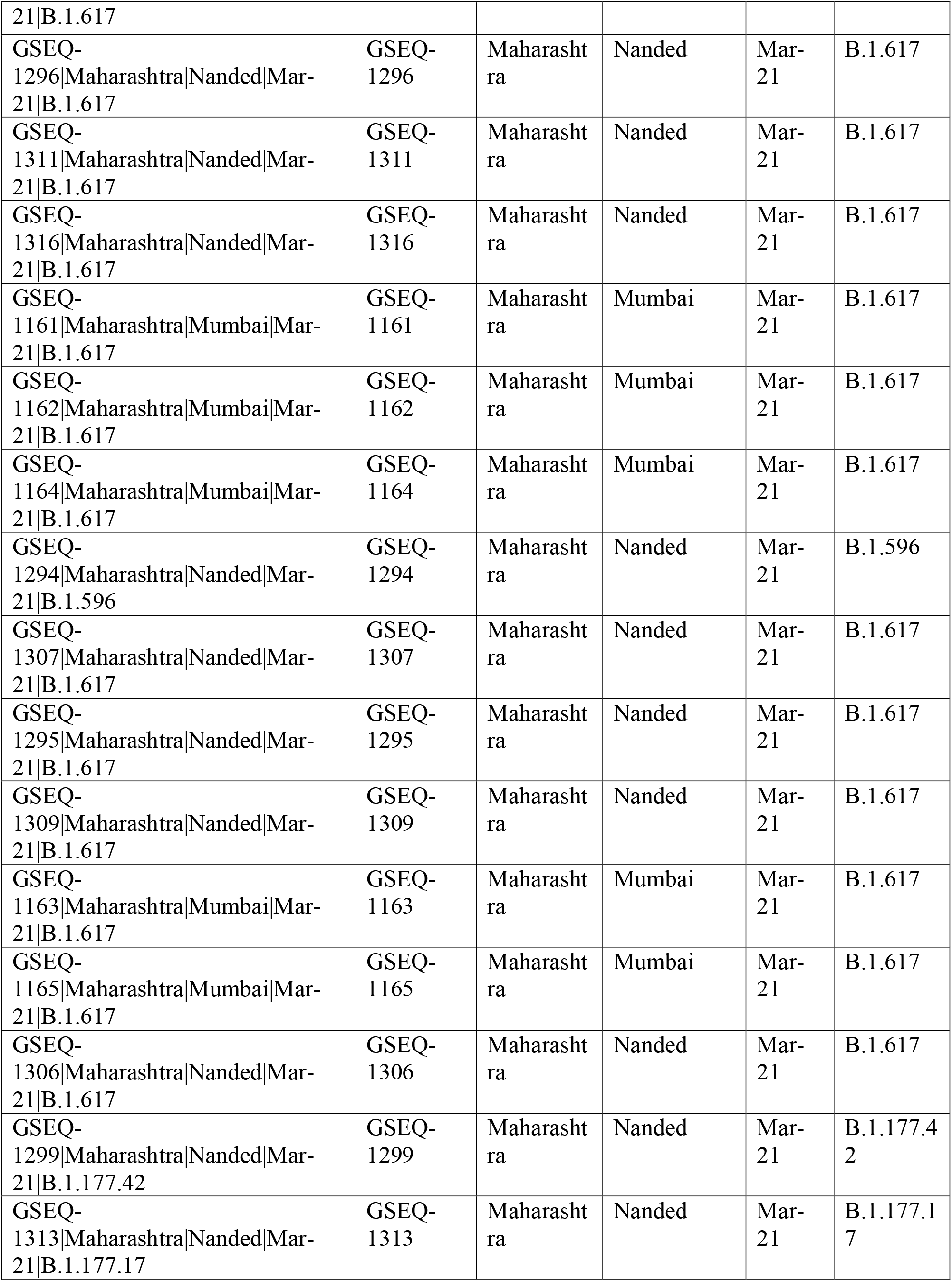
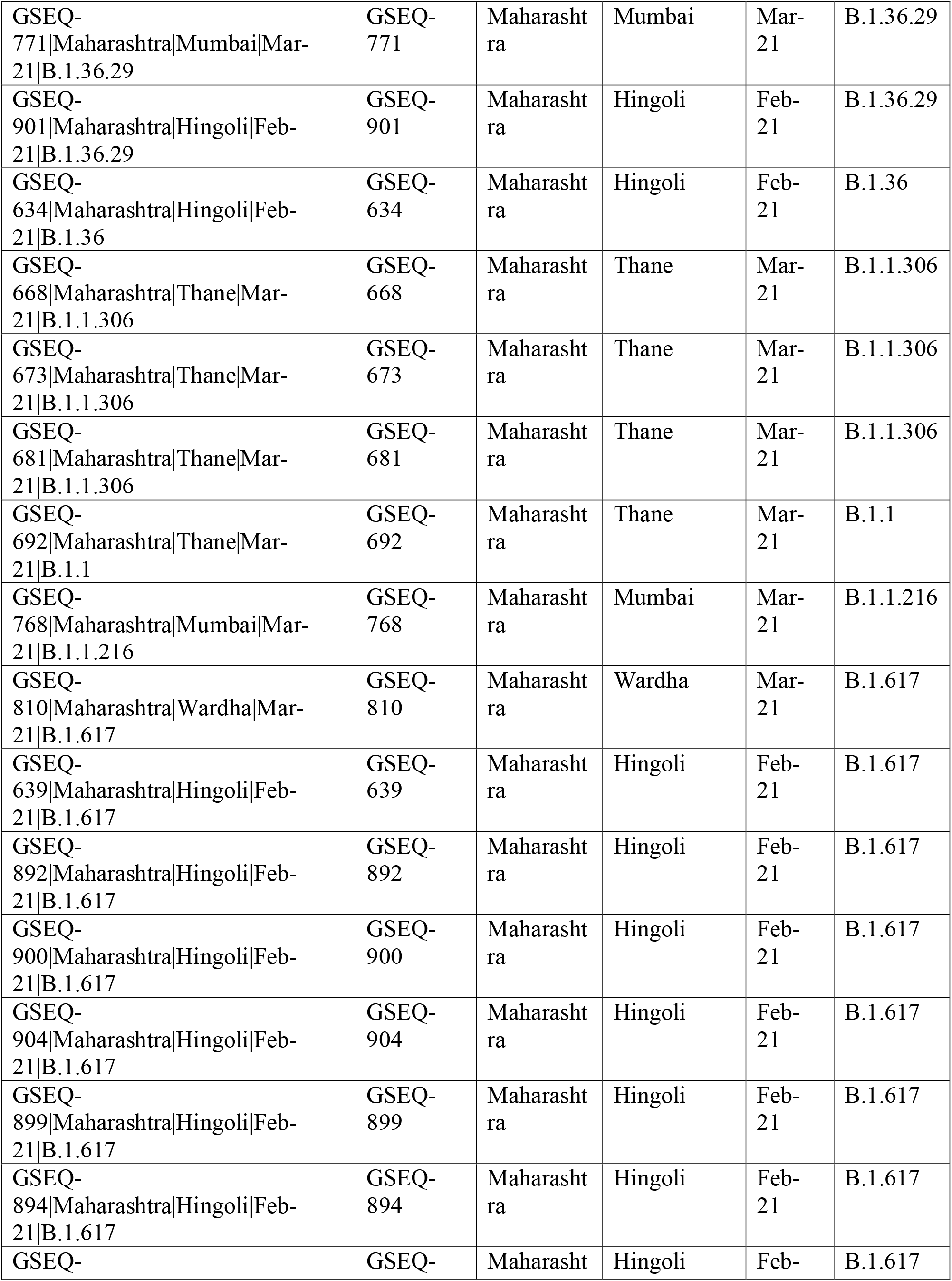
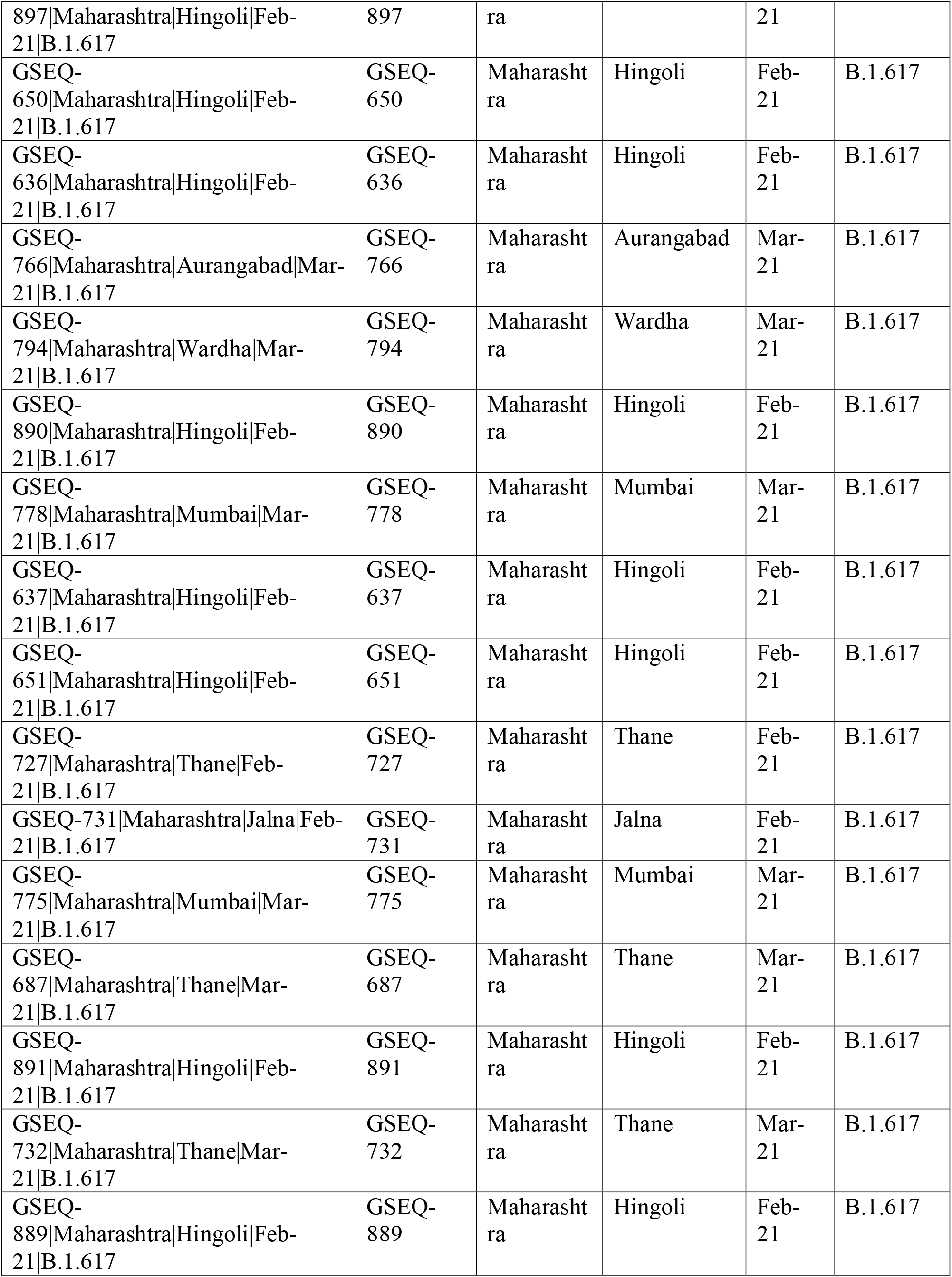
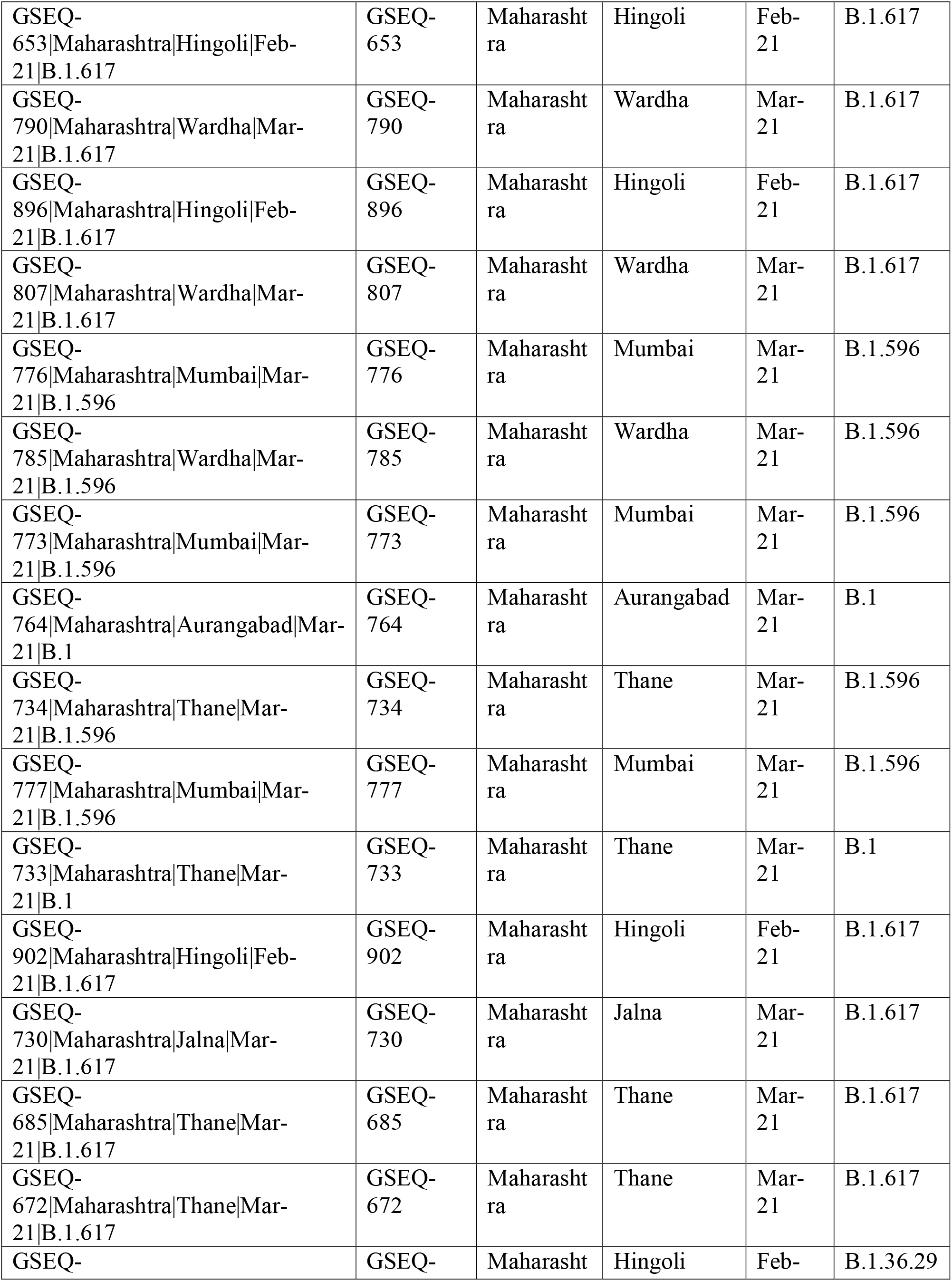
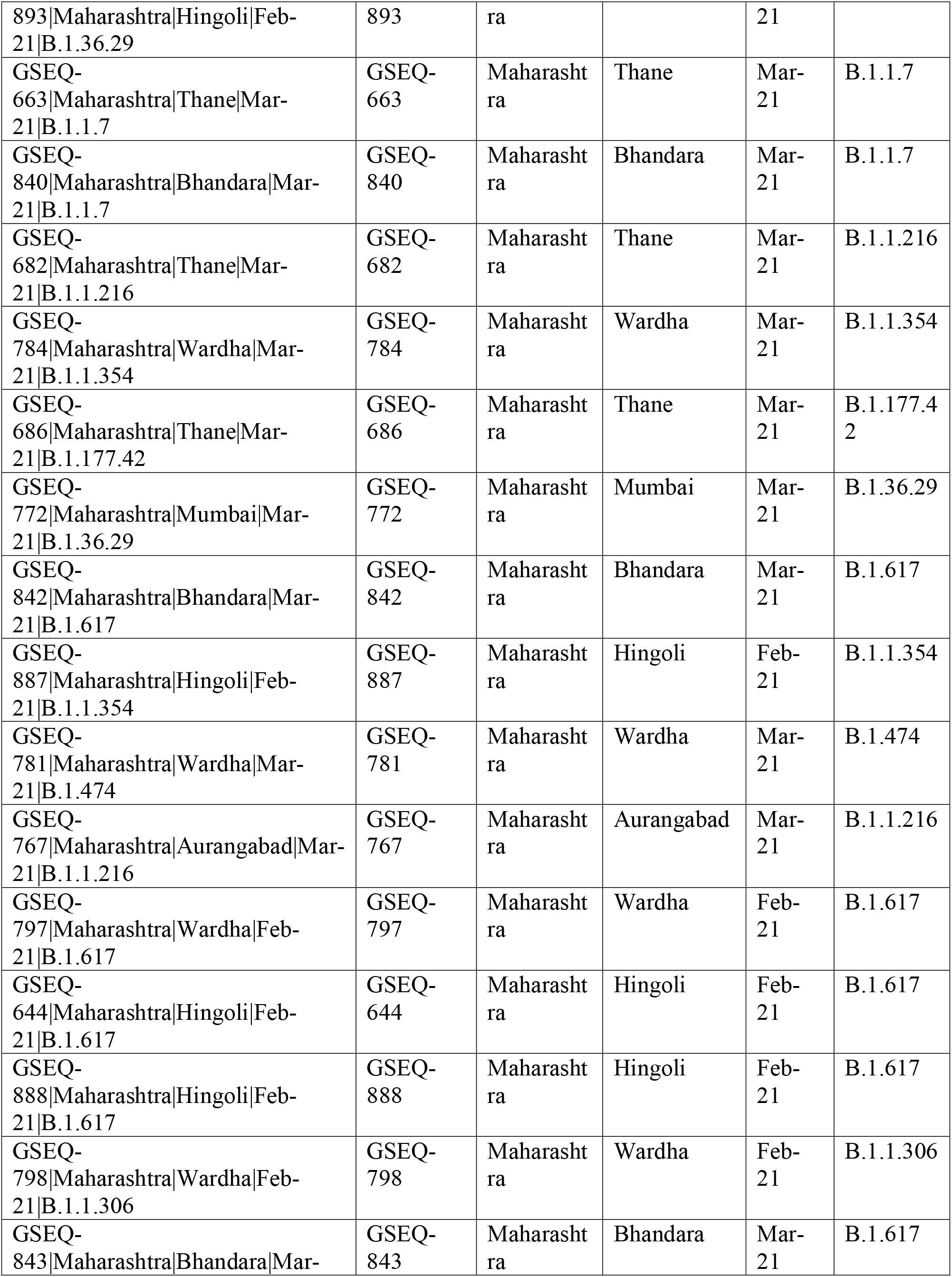
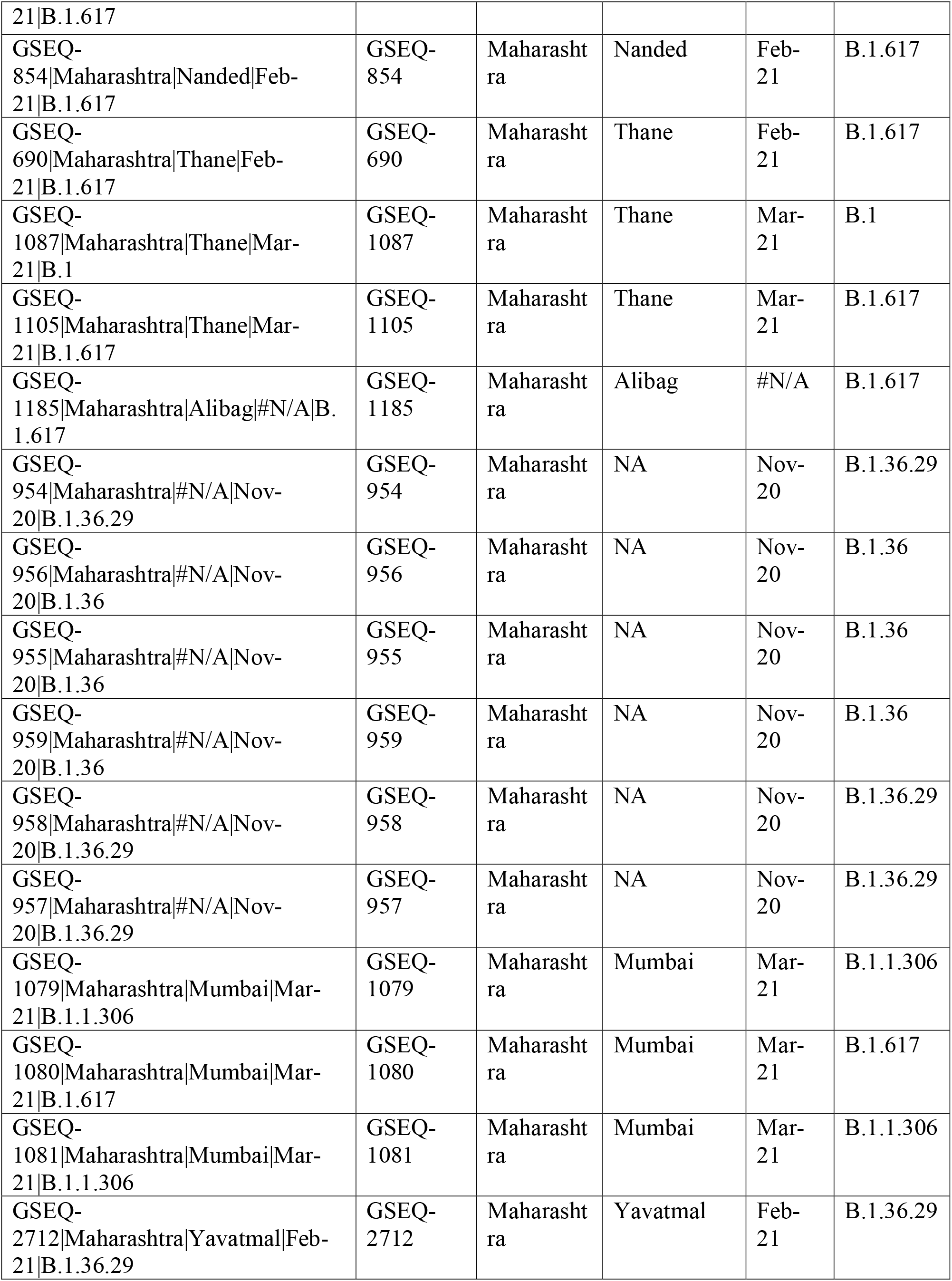
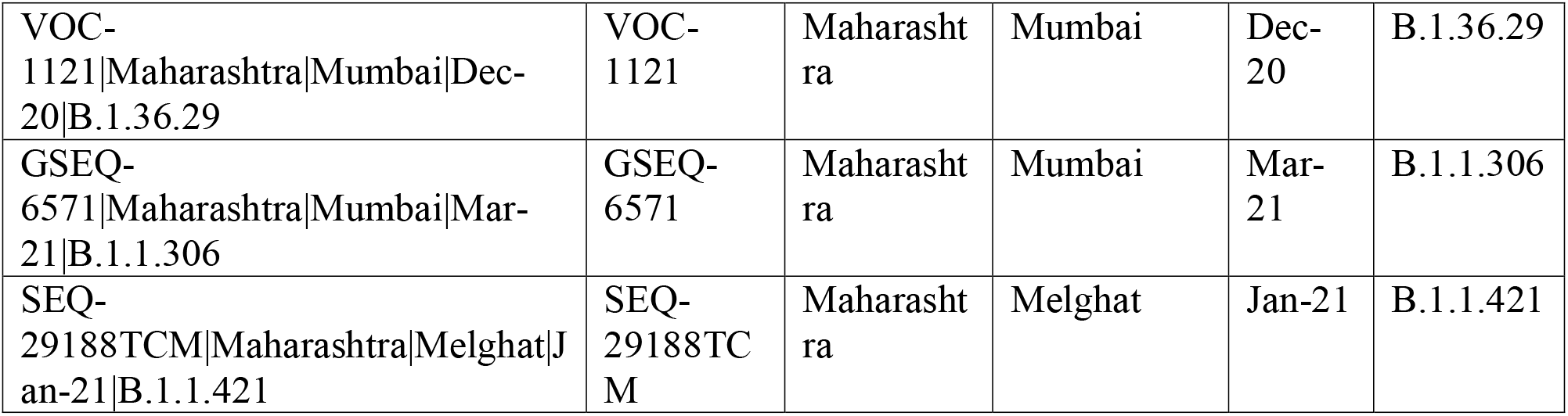
List and details of whole genome sequences used in the study

**Supplementary Table 2:**
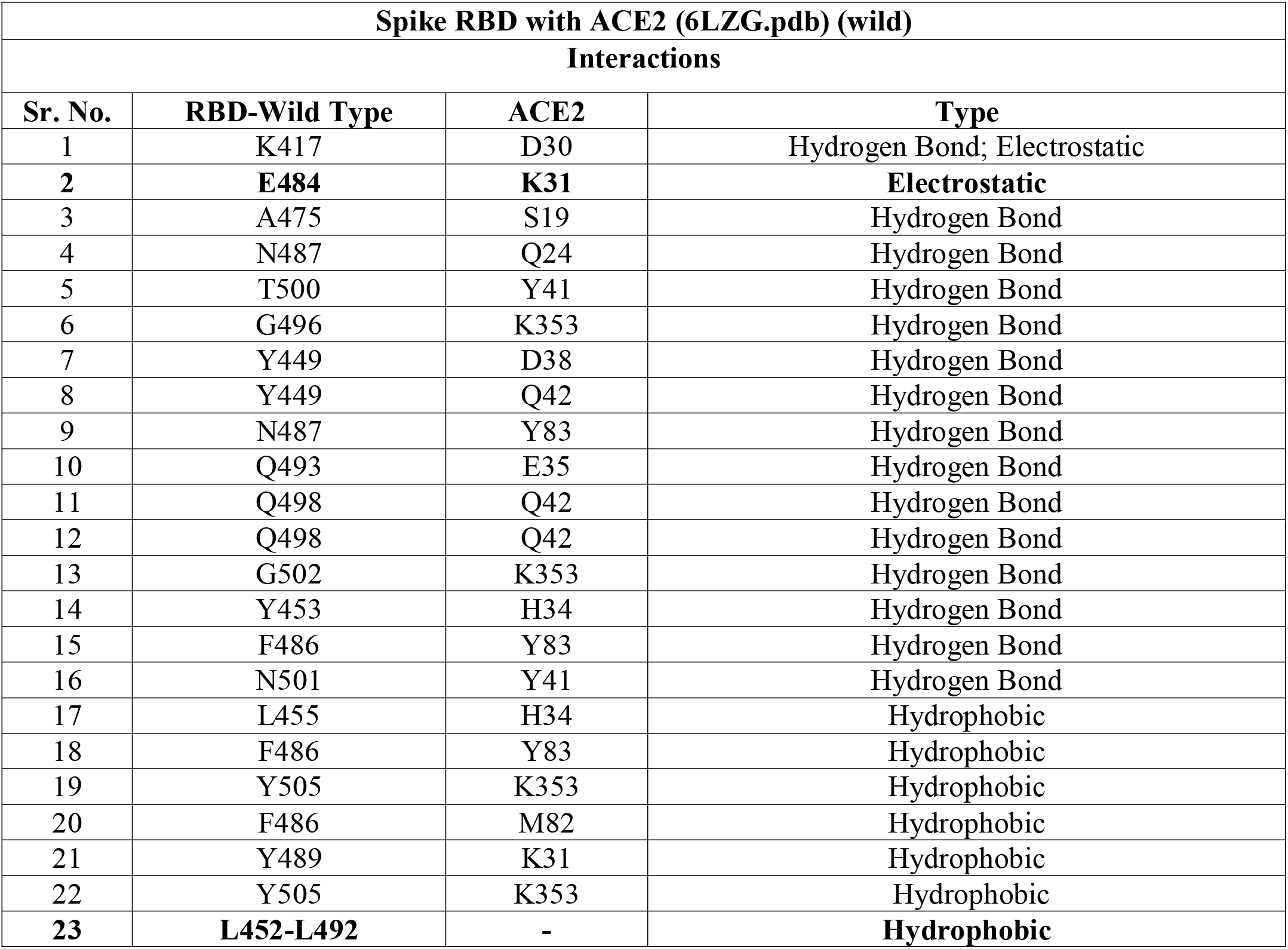
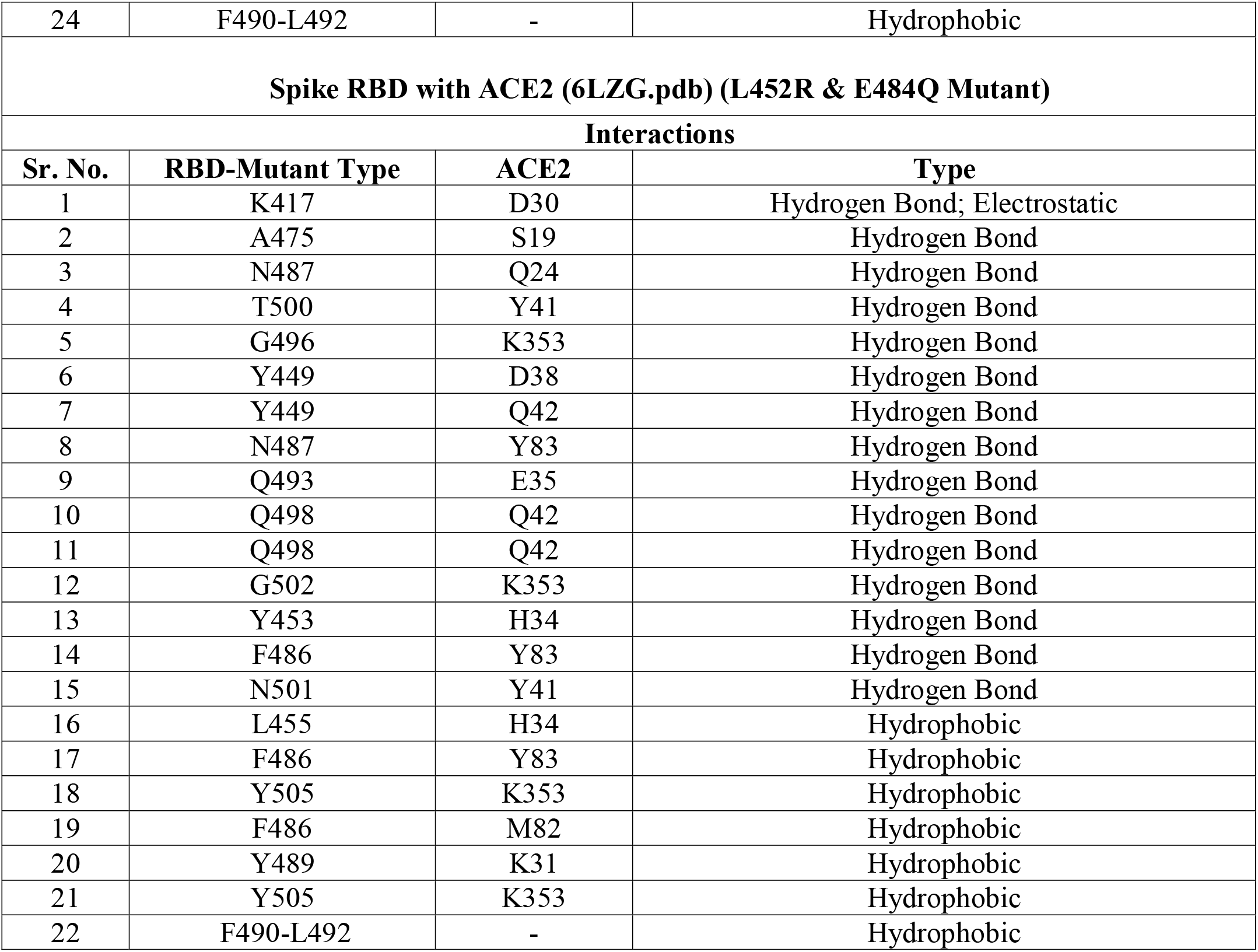
Intra and inter-molecular interactions of wild and mutant spike RBD of SARS-CoV-2 with ACE2. The bonds that are highlighted indicate those that are disrupted in the mutant strain

**Supplementary Table 3:** Intermolecular interaction of neutralizing antibodies with wild and mutant spike RBD of SARS-CoV-2

|  |  |  |  |
| --- | --- | --- | --- |
| A | **Spike RBD with Antibody REGN10933 (Wild)** |  |  |
|  | **Interactions** |  |  |
|  | **RBD** | **Antibody** | **Type** |
| 1 | GLU484 | H: TYR53 | Hydrogen Bond |
|  |  | H:SER56 |  |
| 2 | LYS417 | H:ASP31 | Hydrogen Bond |
|  |  | H:THR28 |  |
| 3 | PHE486 | L: LEU94 | Hydrophobic |
|  |  | H: ARG100 | Electrostatic |
|  |  | H:TYR59 | Hydrogen bond |
| 4 | TYR489 | H: TYR53 | Hydrophobic |
|  | TYR489 | H:TYR33 | Hydrogen Bond |
| 5 | TYR453 | H:ASP31 | Hydrogen Bond |
| 6 | LYS417 | H:ASP31 | Hydrogen Bond |
| 7 | ALA475 | H:MET104 | Hydrophobic bond |
| B | **Spike RBD with Antibody REGN10933 (Mutant)** |  |  |
|  | **Interactions** |  |  |
|  | **RBD** | **Antibody** | **Type** |
| 1 | LYS417 | H:ASP31 | Hydrogen Bond |
|  |  | H: THR28 |  |
| 2 | TYR453 | H :ASP31 | Hydrogen Bond |
| 3 | PHE486 | H: ARG100 | Hydrogen Bond |
|  |  | L: LEU94 |  |
|  |  | H:TYR59 |  |
| 4 | ALA475 | H:MET104 | Hydrophobic bond |
| 5 | TYR489 | H:TYR53 | Hydrophobic |
| 6 | GLN484 | H:SER56 | Hydrogen bond |
| 7 | CYS488 | H:TYR33 | Hydrogen bond |
| C | **Spike RBD with Antibody P2B-2F6(Wild)** |  |  |
|  | **Interactions** |  |  |
|  | **RBD** | **Antibody** | **Type** |
| 1 | LEU452 | H: ILE103 | Hydrophobic |
|  |  | H:VAL105 |  |
| 2 | GLU484 | L: ASN33 | Hydrogen bond |
|  |  | L:TYR34 |  |
| 3 | GLU484 | H:ARG112 | Hydrogen bond; Electrostatic |
| 4 | PHE490 | H: VAL106 | Hydrophobic |
|  |  | H:PRO107 |  |
| 5 | TYR449 | H: SER31 | Hydrogen bond |
| 6 | ASN450 | H: SER30 | Hydrogen bond |
|  |  | H:SER31 |  |
| 7 | GLY447 | H: TYR27 | Hydrogen bond |
| D | **Spike RBD with Antibody P2B-2F6 A(Mutant)** |  |  |
|  | **Interactions** |  |  |
|  | **RBD** | **Antibody** | **Type** |
| 1 | GLN484 | L: ASN33 | Hydrogen bond |
|  |  | L: TYR34 |  |
| 2 | ASN450 | H: SER30 | Hydrogen bond |
|  |  | H:SER31 |  |
| 3 | PHE490 | H: VAL106 | Hydrophobic |
|  |  | H: PRO107 |  |

